# Sensitive, direct detection of non-coding off-target base editor unwinding and editing in primary cells

**DOI:** 10.1101/2025.09.25.678665

**Authors:** Tong Wang, Selin Jessa, Georgi K. Marinov, Sandy Klemm, Anshul Kundaje, William J. Greenleaf

## Abstract

Base editors create precise nucleotide changes in DNA, but their off-target activity remains challenging to quantify. Here, we develop and deploy a direct, *in cellulo* sequencing assay that simultaneously measures both Cas9-mediated unwinding and deaminase editing of genomic DNA (beCasKAS). Our strategy nominates >460-fold more potential off-target sites than other methods by enriching for Cas9-dependent R-loops immediately preceding editing. Using beCasKAS in primary human T-cells, we observe that mRNA-encoded ABE8e and PAMless ABE8e-SpRY base editors have distinct off-target profiles that can be mitigated by optimizing mRNA dose. Finally, we combine beCasKAS with base-resolution deep learning models to risk-stratify off-target edits by their likelihood of epigenetic dysregulation. Collectively, beCasKAS offers a sensitive and facile tool to optimize the balance between base editor on- and off-target activity.

## Main Text

CRISPR base editors are promising genetic editing technologies that underlie at least 19 active clinical trials globally (*1*). Despite the enormous potential of base editing, the impact of off-target activity remains challenging to comprehensively quantify, both because identification of potentially rare edits at off-target sites often requires complex molecular assays of uncertain sensitivity, and because off-target edits impacting non-coding regions are difficult to classify as detrimental or benign. Currently no existing strategy can detect base editor specific off-target effects within primary cells, which would allow a human gene therapy product to meet Food and Drug Administration’s (FDA) ideal recommendations (*2*). Many off-target detection methods are based on *in silico* nomination (*3*, *4*) or *in vitro* biochemical assays (*5–8*), but these approaches generate many false positives and negatives. While *in cellulo* methods exist for detecting off-target base edits, they have not been applied in primary cells likely due to several limitations: these methods optimally detect double-stranded breaks (*9–12*), can only be used on one type of base editor (*13*, *14*), or require whole-genome sequencing (*15*, *16*). As a result, controversy remains on whether off-target base edits are non-detectable, random, or systematically enriched at sites of Cas9 guide RNA (gRNA) binding (*13*, *15*, *16*).

To address the critical need for sensitive detection of CRISPR off-targets, we previously reported CasKAS (**Cas K**ethoxal-**A**ssisted **S**ingle-stranded DNA sequencing), a rapid, inexpensive assay that directly detects DNA editor R-loops genome-wide both *in vitro* and *in cellulo* using the commercially available N_3_-kethoxal reagent (*17*). Although CasKAS can be used to discover the off-target sites made by any ssDNA forming genome editor, we reasoned it would be especially useful for characterizing base editors (Fig. 1A). In most base editors, a C>T or A>G edit can be site-specifically installed by a deaminase within a small single-stranded DNA (ssDNA) editing window created by the Cas9 R-loop (Fig. 1B) (*18*, *19*). The base editing deaminases are intentionally chosen to have only ssDNA and not dsDNA activity to limit off-target toxicity. Furthermore, Cas9 R-loop formation has also been shown to be rate-limiting in biophysical studies (*20*). We therefore posited that an assay detecting R-loop formation, rather than binding, may be especially useful for base editors employing ssDNA deaminases (*21*, *22*). In our base-editor CasKAS (beCasKAS) approach, we use an N_3_-kethoxal reagent to label unwound R-loops, pull these regions down, and use sequencing to detect off-target deaminase edits (Fig. 1C) (*23*).

**Fig. 1:**
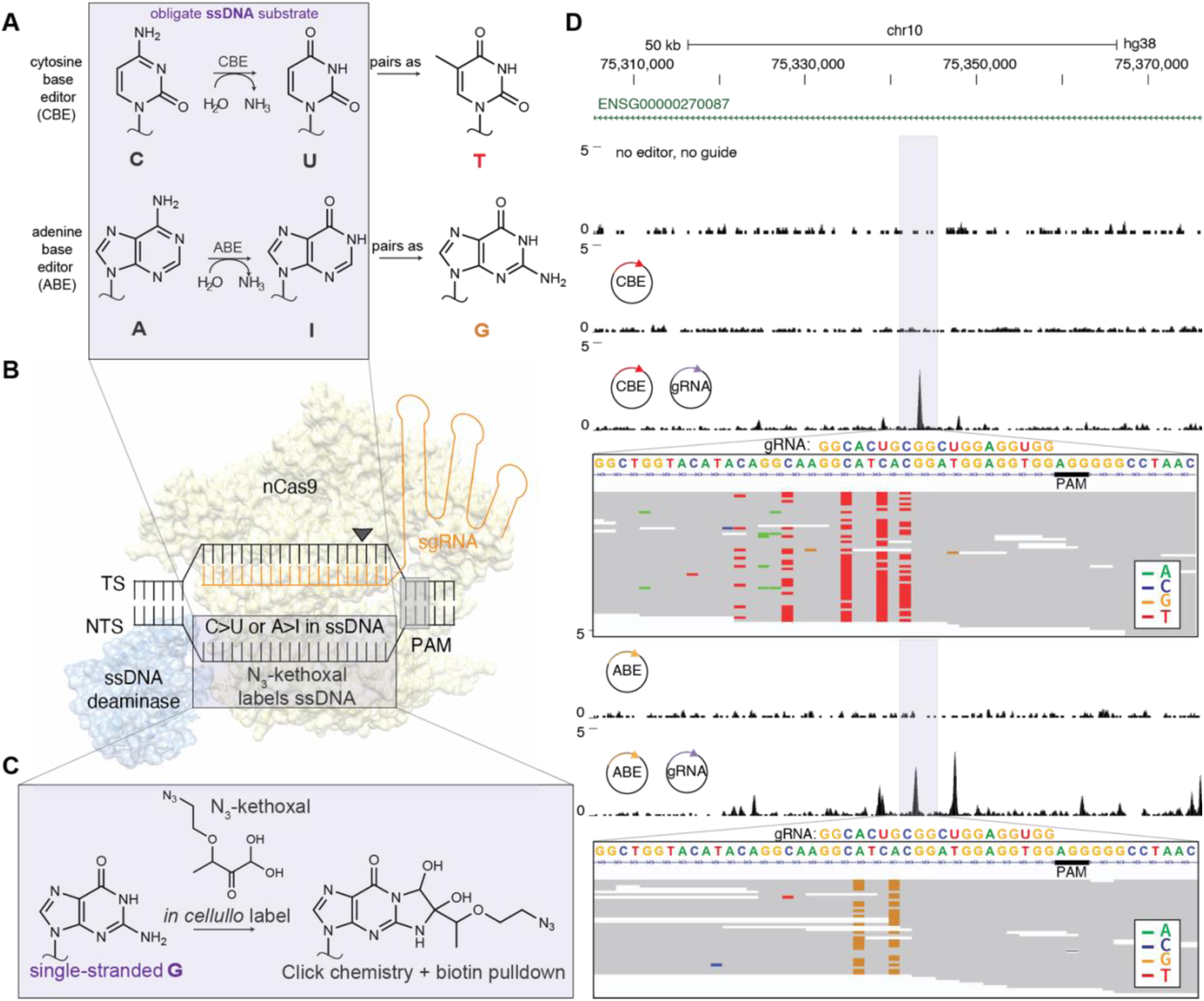
beCasKAS directly detects unwinding and editing *in cellulo.* **A)** Biochemical reactions of CBEs and ABEs. Most CBE and ABE editors require ssDNA substrates. **B)** Cartoon representation of the R-loop intermediate in canonical CBEs and ABEs. CBEs have an additional DNA repair inhibitor that is not shown. **C)** N_3_-kethoxal reagent covalently labels ssDNA at unpaired guanine nucleotides. **D)** Representative genomic track showing a gRNA dependent off-target R-loop. Sequencing reads within the highlighted peak (purple) show C>T edits (red) and G>A edits (green) for the CBE. A>G edits (gold) are seen for the ABE. The targeting gRNA and PAM are shown.

Here, we develop and deploy beCasKAS in both cell lines and primary human T cells. Using beCasKAS we find a wide editing window for the highly active editor eBE and a narrow editing window for the extensively evolved ABE8e (*24*, *25*). We measure deaminase-specific mutational sequence contexts intrinsic to these enzymes and uncover a minimally homologous 5-nucleotide PAM-adjacent seed for unwound R-loops. Surprisingly, we estimate the frequency of gRNA-dependent ABE8e edits in HEK293T cells to be only ∼1.9-fold less than for eBE, although eBE generates ∼20-fold more edits per R-loop than ABE8e. We additionally optimize a primary human T cell compatible beCasKAS workflow and demonstrate a means to quantitatively measure off-targets across increasing concentrations of base editor enzyme, allowing for identification of optimal editor dose for therapeutically relevant clinical targets. Finally, we use a deep learning approach to triage off-target non-coding sites found by beCasKAS predicted to affect DNA accessibility in primary human T-cells.

## Results

### beCasKAS directly detects unwinding and editing in cellulo

We selected the cytosine base editor eBE and adenine base editor ABE8e as two contrasting DNA editors to evaluate by beCasKAS (*24*, *25*). While these editors have never been directly compared experimentally, eBE employs the endogenous human deaminase APOBEC3A (hA3A) which exhibits significant off-target activity compared to other cytosine deaminases and also is known to be dysregulated in multiple human cancers (*26*, *27*). In contrast, ABE8e is a widely used adenine base editor thought to exhibit considerably less off-target activity (*25*). We used the previously studied HEK4 gRNA which allowed us to benchmark against other existing *in cellulo* off-target detection methods (*9*, *11*, *13*).

We transfected base editor and gRNA plasmids into HEK293T cells. After one, two, and three days of editing, cells were treated with the cell-permeable N_3_-kethoxal reagent to covalently label ssDNA. After gDNA extraction, biotin labels were attached to the ssDNA using a click reaction, DNA was sheared, and a streptavidin pulldown was performed before library preparation (see Methods). One day after transfection, gRNA dependent Cas9 R-loops were detected at a highly homologous off-target site (Fig. 1D, purple). At this site, for the CBE, visible C>T (red) and G>A (green, opposite strand C>T) editing was observed near the PAM (gray) distal end of the gRNA sequence. In contrast, only A>G (yellow) edits were observed for the ABE enzyme. Thus, our approach enables detection of both unwound gDNA in R-loops (as gRNA-dependent beCasKAS peaks), as well as individual base edits (as single nucleotide variants in beCasKAS reads).

### beCasKAS quantifies strand-specific behavior of base editors genome-wide

To assess native genome-wide beCasKAS patterns, we measured beCasKAS signal over contiguous, non-overlapping 1-kbp bins after transfection with either ABE or CBE but no gRNA (Fig. 2A). We observed similar ssDNA signals in these two conditions (Pearson’s r = 0.93), although we found four regions which appeared significantly enriched in the CBE-only over ABE-only conditions (Fig. S1A). These reads mapped to exons in the hA3A gene with mismatched bases over intronic regions, suggesting that beCasKAS was detecting transcriptionally active ssDNA from the transfected plasmid and not genomic DNA. We also evaluated the ssDNA landscape with or without the presence of gRNA for the CBE (Fig. 2B). We found that the CBE generated many putative gRNA-dependent R-loops (Pearson’s r = 0.90), and observed strong correlations between beCasKAS replicates (Fig. S1B) performed on the same (Pearson’s r = 0.91) and independent days (Pearson’s r = 0.89).

**Fig. 2:**
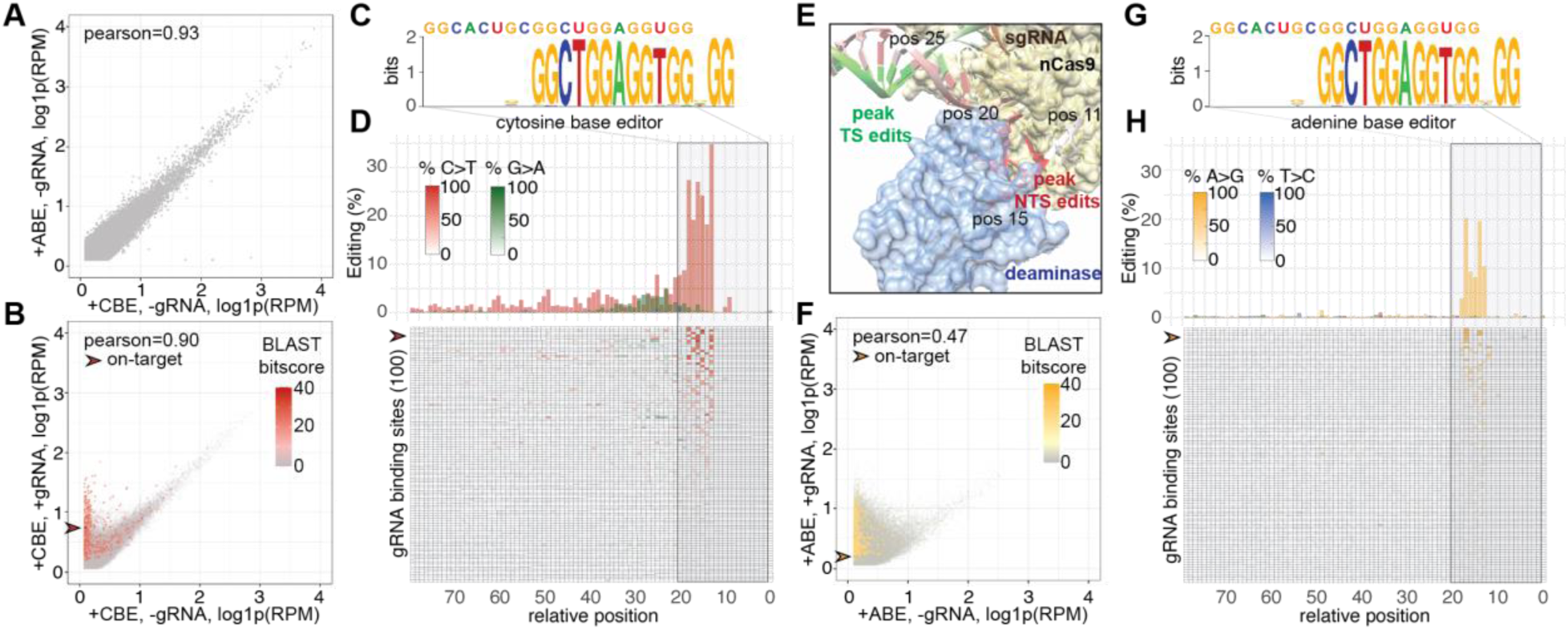
beCasKAS validates known properties of CBE and ABE at sites with significant homology. **A)** Contiguous, non-overlapping 1-kbp bins of genome-wide ssDNA signals detected in beCasKAS without gRNA. **B)** Contiguous, non-overlapping 1-kbp bins for CBE +/- gRNA conditions. Overlaid is the BLAST bitscore. The on-target peak is highlighted with the arrowhead. **C)** All CBE BLAST hits identified after multiple sequence alignment as a WebLogo. **D)** The top 100 most edited BLAST identified regions for CBE are shown as a bar-graph, and each individual site is shown below in heatmap format. Distances are relative to the PAM. The on-target peak is highlighted with the arrowhead. **E)** Editing frequency of cytosine base editor overlaid on a crystal structure of a base editor in complex with DNA (PDB: 6VPC). The numbers correspond to relative position on the gRNA non-target strand (NTS) as in panel **D)**. **F)** Contiguous, non-overlapping 1-kbp bins of ssDNA signals in beCasKAS for ABE +/- gRNA conditions. **G)** All ABE BLAST hits identified after multiple sequence alignment as a WebLogo. **H)** Top-100 most edited BLAST identified regions for ABE.

We next sought to validate highly homologous gRNA off-target sites. We called sharp peaks enriched in pulldown relative to input conditions using MACS2 (henceforth labelled “total peak calls”) and focused on the day 2 time point, which showed significant editing and unwinding of off-target sites (Fig. S1C) (*28*). Following the strategy of previous off-target identification efforts, we used BLAST to determine if any gRNA-homologous regions could be found within the called peaks (*9*, *29*). We found 547 BLAST hits with NGG seeds for the CBE. Among the hits with significant BLAST homology, we further generated a position weight matrix (PWM) to understand the determinants of gRNA-dependent off-target binding (Fig 2C). We found the PAM-adjacent and not PAM-distal bases to be most important, consistent with the existing model for seed-dependent Cas9 interrogation of genomic DNA (*30*, *31*).

We next visualized the top 100 most edited BLAST hits for the CBE and plotted their average conversion frequency as a function of distance from the PAM (Fig. 2D). The on-target site, which was the 4th most edited site overall, showed 68.3%, 78.3%, and 64.4% editing at three target cytosines in positions -18, -16, and -13 relative to the <u>N</u>GG PAM. Across all 100 sites, we found that C>T edits predominantly occur ∼13-19 bases upstream of the PAM, consistent with the intended base editor behavior of converting bases within the editing window of the gRNA non-target strand (NTS). However, NTS edits are also detectable significantly outside of the protospacer, consistent with prior reports using other base editors (*13*, *32*).

We also observed frequent gRNA target strand (TS) edits (G>A changes on the positive strand) that peaked 20-30 bases upstream of the PAM (green), staggered from the C>T edits. Overlaying these editing frequencies on a crystal structure of a base editor in complex with DNA (Fig. 2E) (*33*) provided structural justification of these patterns, as the NTS is most proximal to the deaminase 10-20 bases upstream of the PAM, while the TS is most proximal to the deaminase ∼25 bases upstream of the PAM. We additionally found slight ∼10-11 nucleotide periodicity of C>T edits on the NTS with local maximums occurring at 38, 49, and 60 nucleotides 5’ to the PAM. These findings suggest that the helical turn of DNA affects editing efficiency for out-of-protospacer edits for eBE, and that more broadly, beCasKAS can detect intrinsic strand-specific preferences of base editors.

We repeated this analysis to interrogate genome-wide patterns of beCasKAS applied to ABEs. In contrast to the CBE editor, the ABE dramatically distorted the endogenous ssDNA landscape (Pearson’s r = 0.47. Fig. 2F). Using BLAST, we found 521 candidate off-target sites at the day 2 time-point that were enriched for the PAM-adjacent seed sequence (Fig. 2G). ABE8e does not show any out-of-protospacer or TS edits (blue) and instead only has significant activity in the known target window 13-18 nucleotides upstream of the PAM (Fig 2H) (*25*). The on-target site is also the 4th most edited site, with target A edited at 70%. Taken together, beCasKAS identifies high-probability gRNA-dependent off-target R-loops as these regions recapitulate known, distinct properties of the CBE and ABE tested.

### beCasKAS detects off-target, base-edited R-loops with limited gRNA homology

Although BLAST identifies highly homologous sites enriched by beCasKAS (721 for CBE, 548 for ABE), we observed a large number of regions genome-wide that were not found by BLAST but had differential read counts with and without gRNA (Fig. 2B/F), suggesting that beCasKAS may be identifying *bona fide* off-target sites missed by other approaches (Fig. 3A). To investigate R-loops not identified by BLAST’s homology-directed sequence search (a strategy henceforth labelled “homology” peak calling), we employed a DESeq2-based workflow to identify consensus differentially identified peaks in the +/- gRNA conditions with an FDR of <0.05 (Fig. S1D, a strategy henceforth labelled “differential” peak calling) (*34*). We found that the CBE and ABE enzymes generate 14.9x and 211x, respectively, more differential peaks than found by homology matching for the gRNA sequence (Fig. 3B). This approach nominates 469x more putative off-target sites than the next most sensitive *in cellulo* base editor method (115,355 beCasKAS vs 246 Detect-Seq = 469x), consistent with how beCasKAS enriches for more reaction intermediates preceding uracil generation (*13*). Of the CBE differential peaks, 96.3% were also called in the ABE condition, suggesting these regions are strongly Cas9-dependent (Fig. S1E). In contrast, 91.0% of the gRNA-dependent peaks for the ABE were not identified in the CBE dataset. Because equal amounts of plasmid DNA with the same CMV promoter driving editor expression was delivered for each condition, we speculate that the difference in off-target sites detected in the ABE relative to CBE is likely explained by the ABE’s codon optimization and smaller size (lack of C-terminal uracil DNA glycosylase inhibitor fusion) relative to the CBE.

**Fig. 3:**
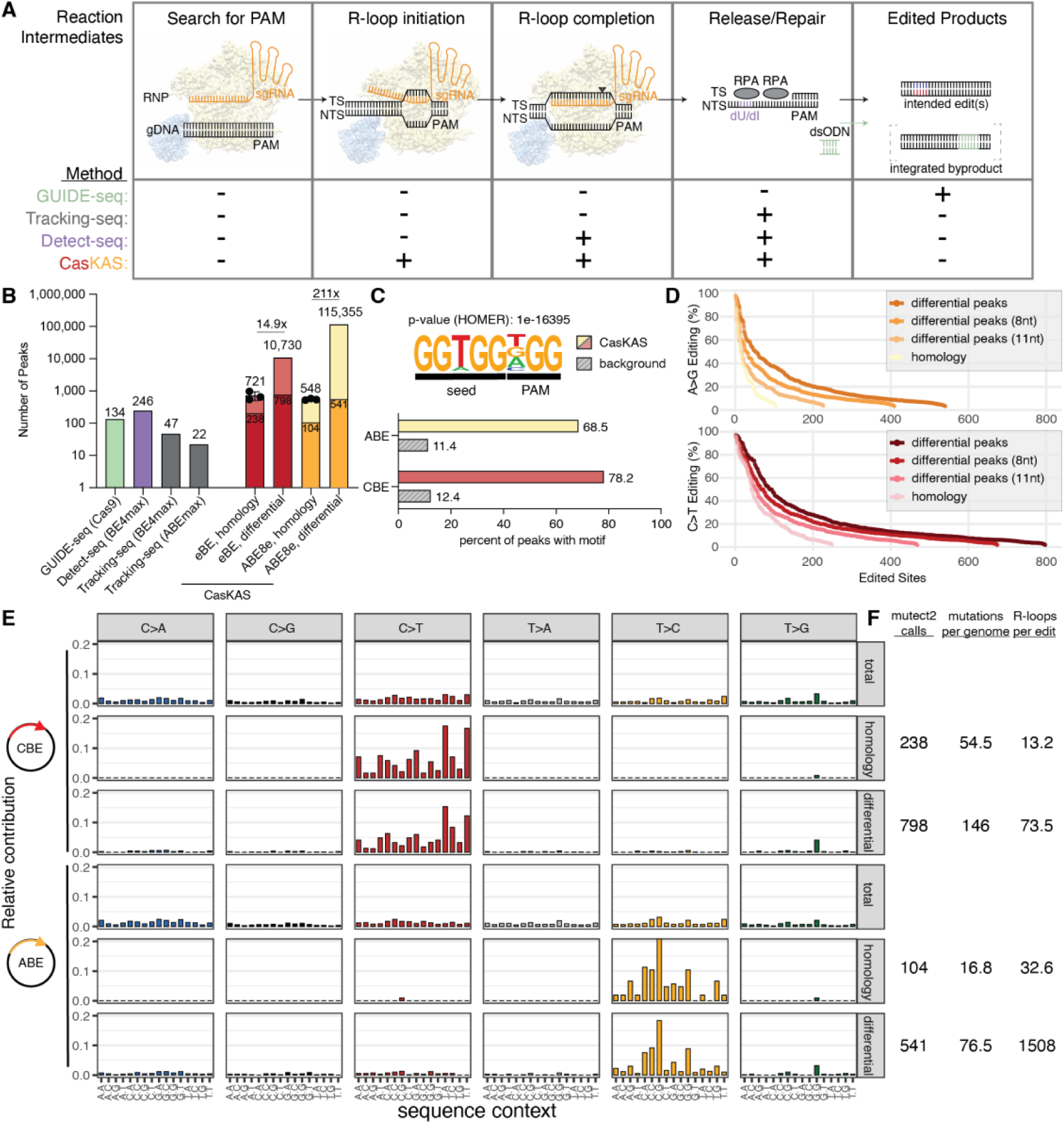
beCasKAS has a highly sensitive, distinct mechanism of enrichment. **A)** Reaction intermediates for base editing. beCasKAS directly measures gRNA-dependent R-loops which occur earlier in time compared to intermediates used by other off-target identification methods. GUIDE-seq measures integration of a transfected oligo (dsODN) at sites with WT Cas9 induced DNA breaks (*11*). Tracking-seq indirectly measures the presence of Replication Protein A (RPA) (*9*). Detect-seq measures deoxyuracil (dU) intermediates generated only by cytosine base editors (*13*). **B)** Comparison of beCasKAS to other *in cellulo* methodologies. beCasKAS R-loops can be found in two ways: homology (BLAST) or differential peaks in BE +/- gRNA conditions (DESeq2 FDR < 0.05 for n=3 conditions). The darker color shows edited peaks. **C)** HOMER *de novo* motif discovery for peaks occurring in differential (DESeq2) peaks versus GC-content matched background. **D)** Rank ordered edited sites identified by differential peaks (DESeq2) and homology (BLAST). “8nt” and “11nt” lines represent R-loops with seed matches of at least that length. **E)** Mutational profile of variant calls. **F)** Absolute editing efficiencies and their relationship to unwound ssDNA peaks for CBE and ABE.

To understand the sequence determinants of the off-target differential peaks, we performed a *de novo* motif search using HOMER (Fig. 3C) (*35*), identifying a 5-nt seed + 3-nt PAM as the top hit for the HEK4 gRNA (p-value: 1e-16395). This motif occurs in 68.5% of putative ABE R-loops, and 78.2% of CBE R-loops while it only occurs in 11.4% or 12.4% of the GC-content matched genomic background, respectively. Thus, gRNA-dependent R-loops are enriched for a 5-nt PAM adjacent seed. Consistent with expectation, we also found that as gRNA seed homology decreases, more off-target sites with detectable base edits are found (Fig. 3D).

To understand the effect of local sequence context on editing efficiency, we next characterized the mutational patterns of both editors. Within the base editor protein, the deaminase enzyme that is chosen contains an evolutionarily divergent recognition loop which confers local sequence context biases to editing efficiency (*36*, *37*). We find that total peak calls (by MACS2) exhibited no mutational sequence preferences, suggesting that the deaminase editing signature is undetectable within regions of genome-wide ssDNA under these assay conditions (Fig. 3E). In contrast, CBE peaks with BLAST homology showed a strong C>T mutational pattern and enrichment of the 5’T<u>C</u> sequence context, the sequence context known to be preferred by the hA3A deaminase (*38*). The 798 total C>T variants in differential peaks also shared the same mutational pattern as the 238 variants observed in homology-based peak identification, suggesting these variants are *bona fide* base editor off-target sites, despite their limited gRNA homology. We also uncovered a consistent ABE8e sequence context mutational pattern for the 104 T>C variants (opposite strand A>G) called in homology peaks and 541 total T>C variants in differential peaks.

We employed amplicon sequencing to investigate how editing within an R-loop is related to absolute editing frequency. We found a linear relationship for both base editors, where the observed editing within the amplicon is 84% and 66% of the editing within the beCasKAS detected R-loop for the CBE and ABE, respectively (Fig. S1F). Using these values as the occupancy of the base editor at every site, we can estimate an absolute mutation frequency of 146 eBE edits per genome and 76.5 ABE8e edits per genome within gRNA dependent R-loops (Fig. 3F). Thus, eBE and ABE8e generate 73.5 and 1,508 peaks per edit, respectively, suggesting that the laboratory-evolved ABE8e deaminase has lower overall ssDNA editing efficiency per R-loop generated than the highly active eBE. Taken together, beCasKAS unearths surprisingly similar absolute mutation frequencies for the promiscuous eBE and the widely deployed ABE8e (146 / 76.5 = 1.91-fold) under these transfection conditions.

### Optimized beCasKAS in primary human T cells

Base editors have recently been used to *ex vivo* engineer allogeneic CAR-T cells for human patients. In one approach, a CD7-targeting CAR-T cell is generated by the simultaneous editing of *TRBC1* and *TRBC2*, *CD52*, and *CD7* to treat T-lymphoblastic leukemia (T-ALL) (*39*). Other genes can be targeted to create allogeneic CAR-T products including *PVR*, *CD3E*, *CIITA*, and *B2M* (*40*). We asked whether beCasKAS data could be used to engineer potentially safer allogeneic products, given these therapies have the potential for widespread clinical application beyond current autologous CAR-T therapies.

We anticipated significant challenges for primary cell beCasKAS as no *in cellulo* base editor specific off-target method currently exists. First, base editor delivery in primary cells cannot be done with plasmids. We instead used 5-methoxyuracil (5moU) modified mRNA (Fig. S2A), a transient delivery method expected to result in fewer off-target edits than plasmid delivery (*41*). Second, activated T-cells divide rapidly, and any cells in S-phase will have exposed ssDNA which will randomly be labelled by our N_3_-kethoxal reagent (Fig. S2B). Finally, electroporation is toxic to T-cells, and extracellular DNA can also be labelled by N_3_-kethoxal. We therefore hypothesized that G1/S-blockade to limit replication, viability sorting, and extracellular nucleic acid degradation with DNase I could all be used to independently improve signal-to-noise in beCasKAS (Fig. S2C). By making these systematic modifications, we optimized the original beCasKAS protocol to improve the on-target signal by 11.1-fold (Fig. S2D). We next sought to compare primary beCasKAS on the same editor (ABE8e) and gRNA (HEK4) as used in HEK293T cells (Fig. 4A). Under these optimal conditions, the read depth-normalized on-target HEK4 gRNA site showed greater peak height relative to background in primary T cells than in HEK293T cells (purple, Fig. 4B).

**Fig. 4:**
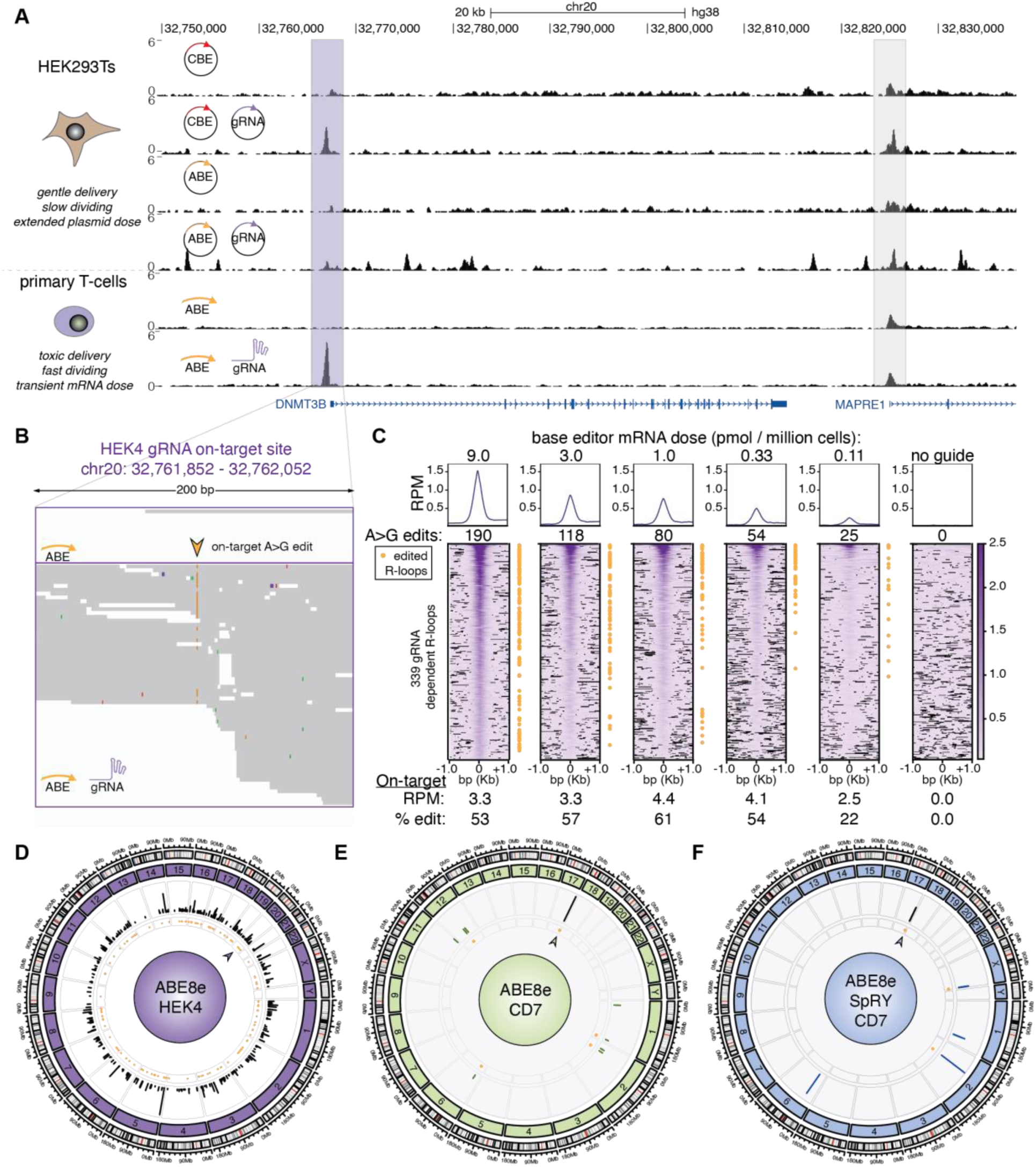
beCasKAS detects off-target sites in primary human cells. **A)** Browser tracks depicting HEK4 gRNA on-target site (purple) and nearby promoter (gray) in HEK293T and primary T cells. **B)** reads at on-target site with target A site marked (arrowhead). **C)** gRNA-dependent R-loops (heatmap) and edits (yellow dot) for the same guide and different base editor mRNA concentrations. **D)** ABE8e and HEK4 gRNA off-target R-loops (lines) and edits (yellow dots). **E)** ABE8e and CD7 gRNA off-targets. **F)** ABE8e-SpRY and CD7 gRNA off-targets. In panels **E)** and **F)**, shared off-target R-loops are shown with a black line while distinct off-targets are shown with colored lines. The arrowhead marks the on-target site in panels **D-F**.

We next aimed to understand whether base editor dose might affect off-target editing. We performed a dilution series with base editor mRNA, while keeping gRNA concentration and cell count constant. While on-target editing efficiency remained constant across a 27-fold dilution series (9.0 to 0.33 pmol mRNA per million T-cells), off-target R-loop formation decreased by 2.8-fold and off-targed editing decreased by 3.5-fold (Fig. 4C, Fig. S4). These results provide proof of principle that systematic mRNA dosing coupled with beCasKAS can provide on- and off-target unwinding and editing measurements that enable identification of an optimal therapeutic dose for base editing.

Although useful for benchmarking the performance of beCasKAS relative to other methods, the HEK4 gRNA is known to be more promiscuous than most therapeutic gRNAs (Fig. 4D). We thus identified off-targets for a splice donor-disrupting CD7 gRNA, which can be used in CD7-targeting CAR-T cells in order to escape fratricide (Fig. 4E) (*39*). In contrast to the 339 R-loops identified with the HEK4 gRNA, the CD7 gRNA identifies only 9 strong candidate R-loops, all with significant seed homology (>10 consecutive bases matching with the protospacer, Fig. S5A). Notably, we identified one off-target edit which creates a missense mutation in the oncogene/tumor suppressor protein NOTCH4 (T428A), with potential bystander edit (H430R), both of undetermined significance (Fig. S5B).

We additionally compared ABE8e to the more versatile, near-PAMless ABE8e-SpRY variant (Fig. 4F) (*42*). Although a PAMless enzyme might be expected to have many off-target sites, we wondered if this would be true using beCasKAS, which uniquely detects R-loop formation (Figure 3A). Across both beCasKAS experiments we found only one shared R-loop, the on-target CD7 peak (A>G edit: 76.3% for WT, 80.0% for SpRY). Interestingly, ABE8e-SpRY had four fewer off-target R-loops (four vs eight) and three fewer edited sites (two versus five) than ABE8e. We also compared R-loops discovered by beCasKAS for PVR/CD3E/CIITA/B2M quadruple edited T-cells. In this setting, the ABE8e-SpRY enzyme exhibited slightly lower editing efficiencies at the four target loci (Fig. S5C). The only shared on-target sites were the four on-target peaks, while we found five unique ABE8e off-target R-loops (three edited) and three unique ABE8e-SpRY off-target R-loops (two edited). Among the off-targets, we identify the known PVR gRNA off-target for the *ICAM-1* exon 2 splice donor, which has been shown to create protein knockdown of *ICAM-1* of uncertain therapeutic significance (Fig. S5D) (*40*). We additionally observed bystander edits which create missense mutations Y110H and W111R which both only occur in reads where the splice donor is also edited. Interestingly, despite having 17 out of 20 matches to the protospacer sequence, using the ABE8e-SpRY enzyme instead of the ABE8e WT enzyme dramatically reduces R-loop formation at this off-target site. Overall, beCasKAS demonstrates that 5moU-modified mRNA delivery of ABE8e or ABE8e-SpRY in primary human T-cells results in limited off-target edits for multiple therapeutic gRNAs. The limited number of R-loops detected for ABE8e-SpRY is also consistent with recent *in vitro* studies on SpRY-Cas9 describing how PAM-relaxed enzymes can be kinetically trapped after initial binding (*43*, *44*).

### Base-resolution deep learning model predicts effects of non-coding off-targets

We next aimed to characterize the consequences of off-target edits for therapeutic gRNAs. Of the 20 off-target R-loops, only three were easily annotated as either missense or splice-disrupting (Fig. 5A). To map the overall *cis*-regulatory element (CRE) landscape and investigate if any off-target sites overlapped active CREs for the 17 intronic and intergenic off-target sites, we performed ATAC-seq in activated primary human T cells. Five sites were found to be in regions of accessible chromatin. Among these five sites, we discovered an ABE8e-SpRY and CD7 gRNA specific off-target site that overlapped with both an accessible region in our ATAC-seq dataset and ENCODE ChIP-seq peaks for the histone marks H3K27Ac and H3K4me1 in activated human T-cells (Fig. 5B).

**Fig. 5:**
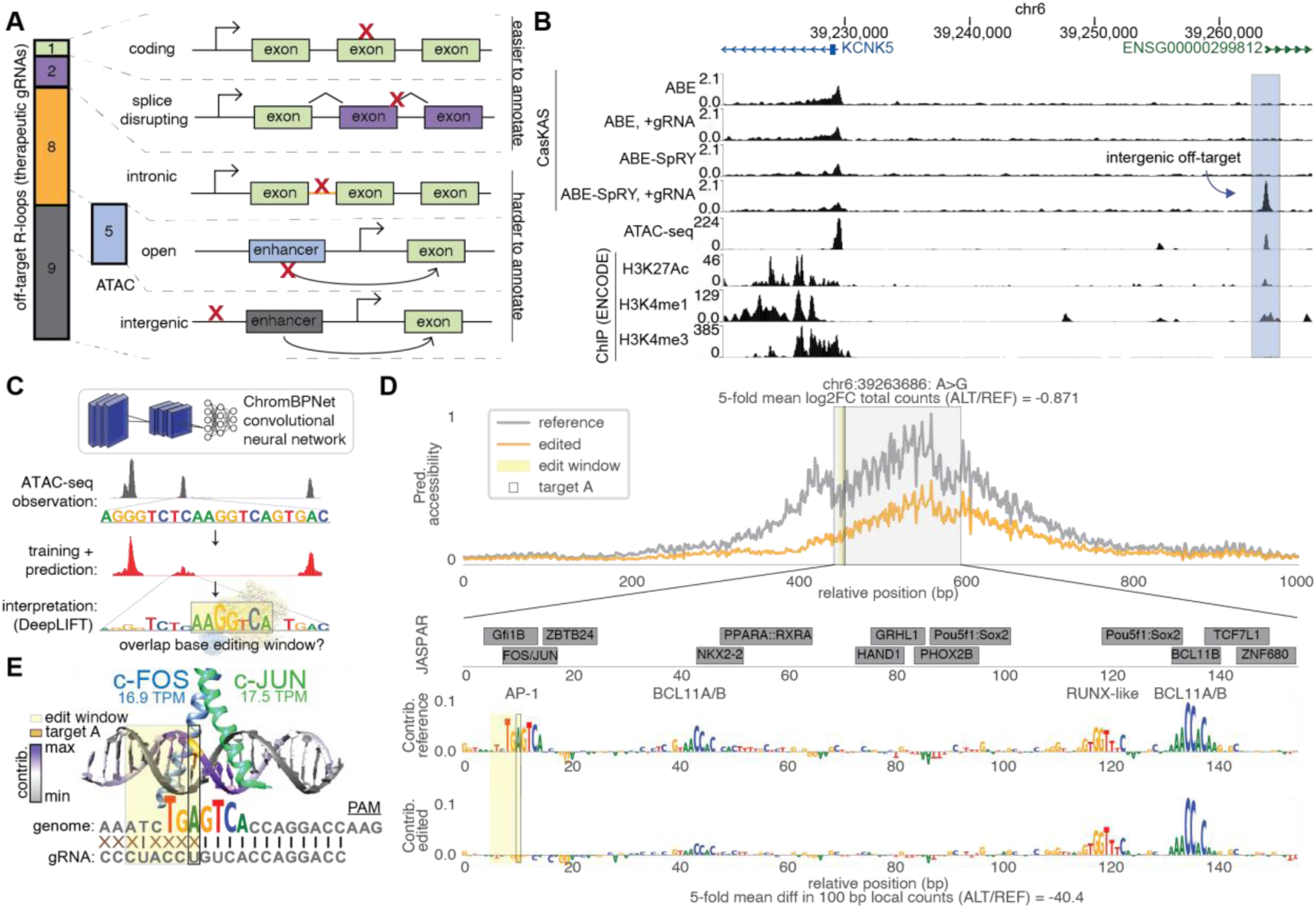
Base-resolution deep learning improves functional annotation of off-target sites overlying a cCRE. **A)** Locations of off-target R-loops for ABE8e and ABE8e-SpRY and five therapeutic gRNAs. **B)** Browser track of novel beCasKAS-detected ABE8e-SpRY + CD7 gRNA off-target site (blue) occurring in a difficult-to-annotate intergenic region, along with ATAC-seq data in primary T-cells generated in this study. The off-target site overlaps with ATAC-seq, H3K27Ac ChIP-seq (ENCODE accession ID ENCFF388LNF), and H3K4me1 ChIP-seq (ENCFF036KBQ), but not H3K4me3 ChIP-seq (ENCFF793LJI). **C)** Overview of deep-learning workflow. A ChromBPNet model is trained on ATAC-seq data to predict accessibility in 1,000 bp windows using 2,114 bp local DNA sequence as input. Models are interpreted by DeepLIFT to derive base-resolution scores representing contribution to accessibility. Editing windows from beCasKAS can be overlaid, with or without an observed mutation. **D)** First track: Predicted accessibility profile at off-target locus for chr6:39263686 A>G edit compared to the reference genome, using T-cell ChromBPNet model. Second track: JASPAR CORE 2024 motifs and base-resolution DeepLIFT contribution scores within the 150-bp ATAC-seq summit. Third track: Contribution scores of unedited (reference) sequence. Last track: Contribution scores after *in silico* chr6:39263686 A>G edit is performed. **E)** Crystal structure of c-Fos/c-Jun:DNA complex (PDB: 1FOS). The A>G edit within the editing window is colored in yellow and the target DNA strand has DeepLIFT contribution scores overlaid. Total RNA-seq quantifications of FOS and JUN expression in activated T-cells is also shown (in TPM units, Transcripts Per Million).

Given that this off-target site could result in a disturbance of an active enhancer element in T-cells, we evaluated the effects of base edits using the ChromBPNet interpretable deep learning framework (Fig. 5C). ChromBPNet models learn to predict chromatin accessibility profiles from ATAC-seq data in 1 kb regions using only the local DNA sequence as input, while learning and correcting for Tn5 transposase bias (*45*). To identify the sequence features driving accessibility, we used the DeepLIFT algorithm (*46*) to interpret the model and extract base-resolution contribution scores in ATAC-seq peaks, representing the influence of each nucleotide on predicted accessibility (Fig. 5C). ChromBPNet has been used to predict the effects of sequence variants on chromatin accessibility, achieving correlations of 0.76 with accessibility effect sizes from molecular quantitative trait loci (QTL) studies, and correlations of 0.67 with expression effect sizes from CRISPR prime editor targeting screens (*45*, *47*). Variants from genome wide association studies (GWAS) prioritized with ChromBPNet are also enriched for statistically fine-mapped variants with high posterior probabilities (*45*). We previously employed this framework to infer the sequence drivers of chromatin accessibility across 189 primary human cell types (*48*).

We first aimed to validate that ChromBPNet models could predict the known therapeutic effect of a well-characterized noncoding variant relevant to therapeutic CRISPR editing. We reasoned that our existing ChromBPNet model trained on human erythroblast ATAC-seq data (*48*) could simulate disruption of the *on-target* enhancer element targeted by the FDA-approved Casgevy (exagamglogene autotemcel). Casgevy is an autologous, *ex vivo* CRISPR-Cas9 gene therapy that disrupts the +58 intronic BCL11A erythroid enhancer in hematopoietic stem cells to restore fetal hemoglobin (HbF) expression for the treatment of sickle cell disease and beta thalassemia (Figure S6A) (*49*, *50*). This +58 intronic BCL11A enhancer is accessible and active only in erythroid cells, and not in other cell types like the T-cells we characterized using ATAC-seq (Figure S6B). ChromBPNet accurately predicted accessibility at the BCL11A enhancers and contribution scores also revealed the known single GATA motif as the primary driver of accessibility at the +58 enhancer (Fig. S6C, middle track). While Casgevy employs CRISPR-Cas9 to disrupt the GATA motif, preclinical base editor studies similarly target this same motif to achieve the same therapeutic phenotype (*51*). We thus simulated the effects on accessibility of two T>C mutations within the ABE8e editing window, using the most optimal gRNA from this study. While one of the T>C edits show minimal effects on overall DNA accessibility (log2FC = -0.222, respectively), the intended on-target base edit results in nearly complete loss of predicted DNA accessibility (log2FC = -1.042) (Figure S6C-D). Base-resolution contribution scores confirmed that only the second T>C edit disrupted the GATA motif, underlying the predicted loss of accessibility (-47.67 change in local counts). Collectively, this experiment shows feasibility of using ChromBPNet to predict changes in accessibility associated with a well-established CRISPR therapeutic target in a non-coding regulatory region.

Given this promising result in the erythroid model, we trained new ChromBPNet models in a five-fold cross-validation scheme on our newly generated, donor-matched T-cell ATAC-seq data, and all models exhibited strong predictive performance (average Pearson’s r = 0.76; Fig. S7A-C, Table S3) For comparison, our prior datasets achieved an average Pearson’s r of 0.78 (IQR [0.75-0.81]; Fig. S7B) (*48*). The contribution scores highlighted four strong motifs corresponding to binding sites for AP-1, BCL11A/B, and RUNX-like transcription factors (TFs) spanning 27 of the 150 bp surrounding the ATAC-seq summit. In contrast, the JASPAR 2024 core database reports at least 13 different motifs spanning 110 of the 150 bp within this same region, demonstrating the difficulty of annotating variants without a cell-type specific deep learning model (*52*). We focused on the AP-1 TG**<u>A</u>**GTCA consensus motif which contained a central A that is both critical for TF target recognition as well as overlies the expected six-nucleotide editing window for ABE8e. The AP-1 family includes dozens of conserved bZIP TFs including the FOS, JUN, and ATF protein families which can be oncogenes or tumor suppressors in different settings, as well as can be intentionally manipulated to overcome CAR-T exhaustion (*53–55*). Consistent with their functional role in T-cells, c-FOS and c-JUN are highly expressed (16.9 TPM and 17.5 TPM, respectively, based on ENCODE RNA-seq data) (*54*). We visualized the well-characterized c-FOS:c-JUN heterodimer interaction with DNA to understand the relationship of the target base, editing window, and DeepLIFT contribution scores (Fig. 5E) (*53*, *56*). At this off-target site, the target A occurs within a G<u>A</u>G sequence context that is the second most favorable for ABE8e editing based on our prior HEK293T experiments (Fig. 3E) and also occurs adjacent to 12 bases of perfect gRNA seed complementarity (Fig. 5E).

We *in silico* installed an A>G mutation within this cCRE and used our ChromBPNet model to predict chromatin accessibility at the edited sequence compared to reference. This single nucleotide change is predicted to significantly decrease peak accessibility (-0.871 log2FC, -40.4-fold change in local counts, Fig. 5D, gold). Moreover, model interpretation revealed that reduced accessibility is driven by complete loss of the impact scores across the entire seven nucleotide AP-1 motif due to this single nucleotide edit. This site has the largest *in silico* predicted effect of all unique off-targets overlapping with regions of open chromatin by ATAC-seq (Fig S7D).

In contrast, we identified example control *in silico* edits which are not expected to create changes to the T-cell epigenetic landscape. For example, mutation of the nearest adenine outside of this core AP-1 motif and within the editing window results in minimal changes in local contribution scores and overall predicted accessibility (0.277 log2FC, Fig. S8A). We also investigated a second independent ABE8e-WT + CD3E gRNA off-target site occurring within a DDIT3:CEBPA JASPAR motif. *In silico* mutagenesis predicts that both As within the core TGC<u>AA</u>T contribute minimally to accessibility at this region (0.066 and 0.033 log2FC, Fig. S8B-D). Instead, a neighboring CTCF motif drives accessibility in this region, consistent with an evident peak in ENCODE ChIP-Seq for CTCF. Thus, this DDIT3:CEBPA edit is not expected to have substantial effects on chromatin accessibility in activated T-cells. Taken together, we show how beCasKAS can uncover base editor specific editing windows that can also be used in concert with interpretable deep learning models trained on epigenetic datasets to triage cell-type specific off-target sites of predicted regulatory importance at single base-pair resolution.

## Discussion

Here, we describe the application of beCasKAS to quantify base editor off-target sites in cell lines and in primary human cells. Unlike other methods that can detect off-targets *in cellulo*, beCasKAS directly quantifies both unwound and edited DNA. Our method only employs commercially available reagents and can be completed within a single day. beCasKAS is especially sensitive for base editor off-targets because R-loop formation must precede ssDNA deaminase editing (Fig 3A, Fig. S9), but our method can in principle be used to probe DNA for any R-loop generating genome editor.

Our results reproduce existing findings for the prevailing model of SpCas9 DNA interrogation (*30*, *31*, *57*). Mechanistically, WT SpCas9 requires a series of concerted conformational changes to catalyze HNH and RuvC nuclease domain activation to create a double-stranded break (*30*). When gRNA mismatches are present, the Cas9 REC2/3 lobes can conformationally prevent nuclease activation (*58*). In contrast, engineered base editors are composed of two tethered proteins without evolved regulation of these conformational checkpoints. As a result, the widely deployed ABE8e exhibits significant potential for mutagenesis, as fully non-complementary substrates can even be unwound and deaminated *in vitro* (*33*). Given this knowledge, our study begs the question of whether detection of an off-target R-loop or a bona-fide off-target edit needs to be seen before a given base editor and gRNA pair are blacklisted. We anticipate the answer will depend heavily on the genomic context. Depending on the setting and base editor used, the relative prevalence of off-target R-loops to off-target edits can range from 74 to 1508-fold (Fig. 3F), although we note that editing is more sensitive to sequencing depth than R-loop formation (Fig. S9). We provide two examples of R-loop formation without clear off-target editing that warrant further investigation: cCREs containing an AP-1 family motif or a Ddit3-CEBPa motif (Fig. 5B-E, Fig. S8).

To explore difficult-to-annotate intergenic off-target sites, we used a cell-type specific deep learning approach to predict the effect of the exact A>G mutations in question on the epigenome. Our results highlight the value of interpretable deep learning strategies for triaging potential off-targets, as our base-resolution model can be easily used to test unintended edits found in human T-cells by any other off-target identification methodology. As models improve and epigenetic measurements grow in size, we anticipate the use of similar approaches to model consequences of non-coding off-target edits on downstream steps of gene regulation including expression. The accurate annotation of non-coding edits will be critical for selecting potent and specific therapeutic gene editors, as non-coding off-target edits could have effect sizes as large as the intentional on-target editing of non-coding regions, such as the BCL11A enhancer (Figure S6D) (*51*).

Our exploration of base editor therapeutic dose is important for three reasons. First, in comparing the mRNA primary T-cell data to the plasmid HEK293T data, we show that the same ABE8e editor + gRNA can result in orders of magnitude change in off-target site generation (115,355 / 339 ∼ 340-fold, Fig. 3B vs 4C), underscoring the importance of FDA recommendations to deploy *in cellulo* assays that accurately recapitulate therapeutic settings. Second, we show that a significant proportion of both editing and unwinding at low-affinity off-target sites can be quantitatively eliminated through careful editor dose choice (Fig. 4C). Finally, dosing a therapeutic candidate at levels exceeding the therapeutic dose might provide a more conservative estimate of an off-target “worse case scenario” for a given model system.

Finally, we note that we deployed beCasKAS to quantify gRNA-dependent off-target sites, rather than potential gRNA-independent off-target sites that may occur due to deaminase activity on ssDNA that exists natively in the nucleus as described by prior studies (*15*, *26*). However, our approach for optimizing primary-cell beCasKAS (Fig. S2) by pharmacologically inducing G1-arrest in activated T-cells might also minimize gRNA-independent off-target editing by limiting ssDNA in replication forks (*59*). We speculate that other pharmacological perturbations might further limit gRNA-independent off-target editing and are actively investigating alternative gene editing workflows with these safeguards in mind.

## Acknowledgments

Cell sorting/flow cytometry analysis for this project was done on instruments in the Stanford Shared FACS Facility (RRID: SCR_017788). We thank Leighton Daigh for suggesting use of palbociclib, Matthew Porteus for critical feedback, and all members of the Greenleaf Lab for helpful conversations.

## Funding

National Institutes of Health grant UM1HG012660 (WJG)

National Institutes of Health grant UM1HG011972 (WJG)

National Institutes of Health grant R01NS128028 (WJG)

National Institutes of Health grant R01HL171611 (WJG)

National Institutes of Health grant DP1HG013599 (WJG)

Arc Institute (WJG)

Chan-Zuckerberg Biohub (WJG)

Wu Tsai Neurosciences Institute Postdoctoral Fellowship (SJ)

CIHR Banting Fellowship (SJ)

## Author contributions

TW conceived of the study. TW performed experiments with assistance from GKM and SK. SJ trained ChromBPNet models and performed in silico mutagenesis with input from AK. GKM performed ATAC-seq. TW and WJG wrote the manuscript, with input from all authors. WJG supervised the work.

## Data and materials availability

FASTQ files will be deposited to GEO upon publication. The following activated primary human T-cell ENCODE datasets were used in this manuscript: CTCF ChIP-seq, ENCFF557ZTZ; H3K27ac Mint-ChIP-seq, ENCSR006QLV; H3K4me1 Mint-ChIP-seq, ENCFF036KBQ; H3K4me3 Mint ChIP-seq, ENCFF793LJI. Code has been deposited at https://github.com/GreenleafLab/ and will be made publicly available upon completion of peer review.

**Supplementary Materials**

Materials and Methods

Figs. S1 to S9

Tables S1 to S3

## Supplementary Materials

## Materials and Methods

### Cell Culture

HEK293T cells were cultured in DMEM + 10% FBS media. All cells were incubated at 37°C with 5% CO2. Primary human CD4+/CD25-T cells were isolated from a leukopak (27 year old female donor, non-smoker, BMI < 30, non-mobilized apheresis collection, consented for full genomic release by STEMCELL Technologies) by negative selection using the manufacturer’s protocol. Cells were stored in liquid nitrogen at 18 million cells per aliquot in 1 mL of 90% FBS with 10% DMSO.

### Cloning

pCMV-hA3A-eBE-Y130F was a gift from Jia Chen (Addgene 113423). Q5 Site Directed Mutagenesis (NEB) was performed as in manufacturer instructions to reintroduce the WT Y130 residue into the eBE plasmid. pCMV-ABE8e was a gift from David Liu (Addgene 138489). pCMV-SpRY-ABE8e was a gift from David Liu (Addgene # 185671). pFYF1548 EMX1 was a gift from Keith Joung (Addgene 47508). Q5 Site Directed Mutagenesis (NEB) was performed to obtain the HEK4 gRNA plasmid. For both ABE8e and ABE8e-SpRY, the coding sequence was PCR amplified from the appropriate plasmid and cloned into a linearized plasmid template with 5’-UTR, 3’-UTR, and poly(A) sequence (Takara 6143). Midipreps (Qiagen) with elution in 10 mM Tris HCl, pH 8.0 were performed to obtain plasmids used in DNA transfection and *in vitro* mRNA transcription. Plasmid integrity was verified by nanopore sequencing (Plasmidsaurus). Plasmids with a poly(A) tail were additionally Sanger sequenced to confirm presence of >100 consecutive adenines (Azenta). Primer and PCR conditions are provided in Table S1.

### HEK293T Transfection

300,000 cells were seeded per well in a 12-well plate the day before transfection to a final volume of 1 mL media per well. Cells were ∼80% confluent the next day. 1.2 μg of base editor plasmid, 400 ng of gRNA plasmid, and 100 ng pMax-GFP plasmid was transfected in each condition. pUC19 plasmid was used to fill total plasmid transfection DNA up to 1.6 μg in conditions without gRNA. 1.6 μg plasmid DNA was diluted in 100 μL Opti-MEM and 6.0 Fugene HD Transfection reagent at 10 min at room temperature before addition to each well. Cells were allowed to grow for 24, 48, or 72 hrs before kethoxal labelling.

### mRNA generation

50 μg of midiprep plasmid DNA was digested in 250 μL reaction volume with rCutSmart Buffer and 20 units of HindIII-HF (NEB) for 1 hr at 37°C. A 0.6x left-sided SPRI bead purification (Beckman) was performed, and DNA was eluted in 20 μL IDTE (10 mM Tris HCl, pH 8.0, 0.1 mM EDTA). DNA was quantified by Qubit 1x dsDNA assay. *In vitro* transcription (IVT) was performed with T7 PrimeCap polymerase (Takara 6144) as in manufacturer instructions with cotranscriptional capping (CleanCap AG, TriLink) and replacing uridine triphosphate with 5-methoxyuridine triphosphate (5moU, TriLink). IVT reactions were incubated at 37°C for 2 hrs with additional brief handmixing after 30 min and 1 hr. 80 U DNase I (Takara) was added and incubated at 37°C for 15 min. Spin columns were used for mRNA purification (NEB Monarch Spin RNA Cleanup Kit) and RNA was eluted in 50 μL IDTE. RNA was quantified by nanodrop and integrity verified by BioAnalyzer (Agilent RNA Nano).

### Primary T-cell electroporation

T cells (∼18 million/mL) were thawed in ImmunoCult basal media (STEMCELL 10981) supplemented with 20% FBS and 20 IU/mL of DNase I (Worthington LS002007), centrifuged at 300 g, and resuspended at 1 million cells/mL in a T75 uncoated flask (Corning) containing complete media comprising basal ImmunoCult media supplemented with recombinant IL-2 (50 IU/mL), Pen/Strep (100 U/mL), and CD3/CD28/CD2 activator (STEMCELL 10970). T-cell media always contained IL2 and Pen/Strep at the same concentrations in subsequent steps. On day 3 after thawing, cells were diluted in T-cell media to a concentration of 150,000 cells / mL. On day 5, 1 million cells were diluted in 20 μL of electroporation buffer (Lonza, 90% P3, 10% Supplement), 1 μL 5moU-modified base editor mRNA (1 pmol / μL in IDTE, except for dilution series), and 1 μL sgRNA (30 pmol / μL in IDTE, IDT Alt-R™, standard desalting, RUO-grade, Table S2) before electroporation with a Lonza 4D-Nucleofector and pulse code EO-115. Cells were recovered for 5 min at room temperature before addition of 100 μL T-cell media per cuvette. Cells from 4 cuvettes were pooled (4 million cells electroporated per condition) into single wells of a 12-well plate and diluted to a 1.6 mL final volume per well. After 3 hrs, cells were G1 arrested with 5 μM palbociclib (Selleck) before allowing to grow overnight for ∼20 hrs before further processing (*60*).

### beCasKAS experiments

beCasKAS experiments were carried out as previously described (*17*) with some modifications. 1 μL of 500 mM N_3_-kethoxal (APE-Bio, dissolved in 100% DMSO) was diluted to 5 mM in 100 μL media per sample and pre-heated to 37°C. For HEK293T cells, media was removed from cells and a PBS wash was performed. Cells were trypsinized, and trypsin was quenched with media before transfer to 1.7-mL microfuge tubes. Cells were pelleted for 5 min at 500 g and resuspended in 100 μL of 5 mM N_3_-kethoxal, and cells were labeled at 37°C in a Thermomixer at 500 rpm for 10 min.

For primary T cells, 20 U DNase I (NEB) and 0.8 U Proteinase K (NEB) were added to the media before harvesting, and cells were placed back in the incubator for 30 min at 37°C. Cells were transferred to 1.7 mL microfuge tubes. Cells were pelleted for 5 min at 500 g and washed once with PBS. Cells were resuspended in Stain Buffer with FBS (BD Biosciences) and DAPI (final concentration: 1 ug/mL). Cells were strained with a 35 μm mesh cap and collected in polystyrene test tubes (Fisher) before flow sorting for DAPI-negative single cells (FACSAria Fusion, Fig. S3). Cells were pelleted for 5 min at 500 g and the supernatant was removed. The cells were then resuspended in 100 μL of 5 mM N_3_-kethoxal and labeled at 37°C and 500 rpm for 10 min in a Thermomixer.

Genomic DNA was immediately obtained using spin column purification (NEB Monarch) and eluted in 100 μL 25 mM Sodium Borate, pH 7 (Teknova) with 10 mM EDTA. gDNA was quantified by Qubit 1x dsDNA reagent. Click reaction was carried out using 87.5 μL of purified gDNA (5-15 ng/μL), 2.5 μL 20 mM DBCO-PEG4 (Sigma) and 10 μL 10x PBS, incubated at 37°C and 500 rpm for 90 min in a Thermomixer. DNA was then purified again using a spin column (NEB Monarch) and eluted in 130 μL 25 mM Sodium Borate, pH 7 without EDTA. DNA was then sheared down to 150-400 bp using a Covaris E220 (peak incident power 175 W, duty factor 10%, cycles per burst 200, treatment time 200 seconds). For biotin pull down, 10 μL of Dynabeads MyOne Streptavidin C1 beads (Thermo) were used per sample. Beads were first washed with 1x B&W before resuspension in 120 μL 2x B&W (10 mM Tris HCl pH 7.5, 1 mM EDTA, 2 M NaCl, 0.1% Tween-20). A portion (10 μL) of sheared samples was saved as input. The remaining 120 μL of sheared DNA were added to the resuspended beads and incubated for 15 min with gentle rotation at room temperature. The samples were placed on a magnet and the supernatant was removed. DNA was washed a total of four times with 1x B&W. During each wash, the sample was incubated in a Thermomixer at 55°C and 1000 rpm for 1 min before placing it on a magnet rack and removing the supernatant. DNA was eluted by adding 16 μL nuclease-free H_2_O to each sample (and 6 μL H_2_O to input samples) and heated to 95°C for 10 min. 15 μL used as input to library preparation. Libraries were prepared using the IDT xGen™ Methyl-Seq Lib Prep kit (10009860) following all steps for <10 ng input gDNA manufacturer instructions (17 PCR cycles). Libraries were quantified using the Qubit 1x dsDNA kit and fragment distribution visualized by Tapestation before pooling to 4 nM in IDTE buffer (Fig. S2E).

Libraries were sequenced using a NextSeq 550 High Output kit (in 2 x 150 bp format) following manufacturer instructions or through Novogene using one lane of a NovaSeq X 10B kit (2 x 150 bp) with a goal of obtaining 20-40 million reads per library. We note that 2x150-bp sequencing is often necessary to capture the middle of the DNA fragment containing the Cas9 binding site, although more stringent shearing and size selection could be used if shorter sequencing reads are desired.

### Cell cycle analysis

T-cells were thawed and activated as described above. On day 3 after activation, cells were diluted to 150,000 cells / mL and 500 μL was plated in individual wells of a 24-well plate. On day 4, palbociclib or equivalent concentration of DMSO was added to the media and cells were blocked overnight for ∼20 hours. The next day, BrdU and 7-AAD based cell cycle labelling was performed according to manufacturer instructions (BD Pharmingen 559619) with a 2-hour BrdU pulse (Fig. S2B).

### Amplicon sequencing

Twelve amplicons were designed using the IDT rhAmpSeq software, and rhAmpSeq libraries were generated according to manufacturer instructions. Libraries were quantified using the Qubit 1x dsDNA kit and fragment distribution was evaluated using a Tapestation before pooling to 4 nM in IDTE buffer. Amplicons were analyzed using CRISPResso2 (*61*). The correlation of the R-loop allele fraction from beCasKAS and absolute allele fraction from each amplicon was calculated to compute a best fit line by linear regression, where the slope represents the average occupancy of the base editor at each binding site. Absolute mutation frequency (per genome) is computed by summing the allele fraction (*AF*) at *n* edited sites identified by DESeq2.

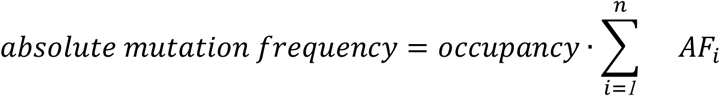

### KAS-seq data processing

Sequencing reads were trimmed using Trim Galore! specifically removing 12 bases from the 5’ end of R2 per IDT instructions. The KAS-Analyzer wrapper was used for subsequent standard steps including alignment to the hg38 version of the human genome using Bowtie2 (*62*, *63*). Reads were deduplicated using gencore (https://github.com/OpenGene/gencore) which allows for consensus duplicates to be obtained. Signal was normalized to Reads per million (RPM) to make samples sequenced to different read depths directly comparable.

The deeptools multiBigwigSummary and plotCorrelation functions were used to visualize global beCasKAS signals (*64*). Point-source sharp peaks were called using MACS2 against an Input control (q-value = 0.01) (*28*). BLAST peaks were called using the blastn-short algorithm and the gRNA sequence as the query (*29*). Clustal Omega was used for multiple sequence alignment to build a position weight matrix (*65*). The aligned nucleotides could be further processed with perbase (https://github.com/sstadick/perbase) to visualize strand-specific editing of each base editor. Protein structures are visualized in UCSF ChimeraX (*66*).

Differential peaks +/-gRNA are called using the DiffBind package using the DESeq2 algorithm (FDR < 0.05, FC > 1) and only the summit (middle 100 bp) was used to search for subsequent variants (*34*). *De novo* motif search was performed using HOMER on differential peaks (*35*). The GATK mutect2 function was used to call variants in “tumor-only” mode (*67*). Variants were atomized with bcftools and any variants aligning to alternate chromosomes were removed. For HEK293T cells experiments, previously identified somatic variants from the HEK293T cell line were removed (*68*). Variants were visualized using the MutationalPatterns R package (*69*).

For the primary T-cell mRNA dilution series, the deeptools computeMatrix and plotHeatmap functions were used to visualize gRNA-dependent R-loops generated by the highest concentration of editor (9 pmol / million). Circos plots were generated using the circlize R package (*70*).

### Primary T-cell ATAC-seq

Cells were thawed and activated in the same way as for primary T cell beCasKAS. On day 3 after activation, cells were G1-blocked with 5 μM palbociclib (Selleck) before allowing them to grow overnight for ∼20 hrs. The Fast-ATAC protocol was used as previously described (*71*). Briefly, cells (50-200,000) were harvested by centrifugation at 500 g for 5 minutes, and the supernatant was removed, then resuspended in 50 μl of transposase mixture (25 μl of 2× TD buffer, 2.5 μl of TDE1, 0.5 μl of 1% digitonin, and 22 μl of nuclease-free water), and incubated at 37°C for 30 min in a ThermoMixer with agitation at 1,000 rpm. The reaction was immediately stopped by adding 250 μl of PB Buffer, and DNA was purified using the MinElute Reaction Cleanup kit (QIAGEN), eluting in 20 μl of Elution Buffer. Final libraries were generated as previously described (*72*): 20 μl of DNA, 2.5 μl of each of the two indexed primers, 25 μl 2x NEBNext High-Fidelity PCR Master Mix, with an initial extension and fill-in for 5 minutes at 72°C; followed by initial denaturation for 30 sec at 98°C, and 10 cycles of 10 sec at 98°C, 30 sec at 63°C, and 30 sec at 72°C. Libraries were purified using the Qiagen MinElute kit, quantified using Qubit 1x dsDNA kit, evaluated on a TapeStation, and then sequenced on an Illumina NextSeq 550 as 2 × 38mers.

### ATAC-seq data processing

Computational processing was carried out as previously described (*73*). Demultiplexed FASTQ files were mapped to the GRCh38 (hg38) assembly of the human genome as 2 × 36mers using Bowtie with the settings “-v 2 -k 2 -m 1 --best --strata” (*74*). Mitochondrial mapping reads were filtered out, and duplicate reads were removed using picard-tools MarkDuplicates. Reads were also mapped separately to the mitochondrial genome using the Bowtie settings “-v 2 -a --best -- strata”, to estimate the extent of mitochondrial contamination. Three technical replicates were pooled for subsequent ChromBPNet model training.

### ChromBPNet model training and interpretation

ChromBPNet models are supervised convolutional neural networks trained to use 2,114-bp one hot-encoded DNA sequence in peaks and background regions to predict the accessibility profile (as a probability distribution) and total natural log counts (as a scalar value) in the central 1,000-bp window of input regions (*45*).

To define genomic regions for training ChromBPNet models, we followed our prior workflow. First, we defined a lenient set of accessible regions. Using our T-cell ATAC-seq data, we first derived pseudoreplicates. For each ATAC-seq fragment, starts and ends (corresponding to Tn5 insertion sites) were randomly allocated to each of two pseudoreplicate files, and pseudoreplicate files were also concatenated into a total-pseudoreplicate file. Macs2 (v2.2.9.1) was used to call peaks on all three pseudoreplicate files with parameters: *-p 0.01 --shift -75 --extsize 150 --nomodel-B --SPMR --keep-dup all --call-summits*. Only peaks called on the total-pseudoreplicate which overlapped peaks called in both pseudoreplicates were retained. Peaks overlapping the GRCh38 ENCODE blacklist (ENCODE accession ENCFF356LFX) were excluded. Peak coordinates were adjusted to 1,000 bp centered at the Macs2 peak summit. Pseudoreplicates were only used for peak calling, and pseudobulk fragment files were used for downstream model training.

We used the ChromBPNet package (https://github.com/kundajelab/chrombpnet, commit a5c231) and followed the workflow described by Pampari et al (*45*). We used the command *chrombpnet prep nonpeaks* to define background regions which match the GC content of peak regions. For each cell type, we used a five-fold cross-validation scheme, where each fold (designated 0 to 4) comprised a different set of training, validation, and test chromosomes, with each chromosome in the test set of at least one fold. We used the default human chromosome folds provided with ChromBPNet (https://doi.org/10.5281/ZENODO.7445373).

ChromBPNet models use a pre-trained bias model and explain the residual accessibility not captured by Tn5 enzyme bias. We trained a bias model to learn the enzymatic bias in our ATAC-seq setting using the fold 0 chromosome split, with bias threshold factor -b 0.9 using the chrombpnet bias pipeline, which also performs model interpretation using DeepLIFT (*45*, *46*). We confirmed that the bias model learned the Tn5 motifs but not transcription factor motifs, and used this bias model to subsequently train ChromBPNet models using the chrombpnet pipeline command with the GRCh38 reference genome from ENCODE. Models were evaluated based on the Pearson and Spearman correlations between predicted and observed log counts in peaks and the Jensen-Shannon Distance between predicted and observed profiles in peaks, for peaks on held-out test-set chromosomes (Fig. S7A-B, Table S3). To generate the average predicted accessibility tracks across folds for peak regions (representing counts per base), for each region, the mean predicted profile logits across folds were softmaxed to convert them to probabilities, then scaled by the exponentiated mean predicted log counts across folds.

We performed model interpretation to determine the extent to which each nucleotide was predictive for accessibility. We ran the chrombpnet interpret command which uses the DeepLIFT algorithm to compute contribution scores for each nucleotide in the 2,114 bp input windows with respect to the predicted counts. Contribution scores were derived for each model fold for all peak regions, and the mean computed across folds. The averaged predicted accessibility profiles and contribution scores were converted to bigWig files for visualization in genome browsers, as well as used for all analyses and figures.

### Prediction of variant effect using ChromBPNet models

We predicted and interpreted effects of specific noncoding variants on chromatin accessibility using trained ChromBPNet models, as we have done previously (*48*). We used the tangermeme package (v0.4.3) for predictions and model interpretation (*75*). For each variant, we used the 1,000 bp model training peaks, and extracted the reference genome sequence for the peak which the variant overlapped. This sequence (extended equally on either side to 2,114 bp) was fed to all five fold trained T-cell ChromBPNet models, to obtain predicted accessibility profile and aggregate log counts in the peak. For each fold, to transform predicted profile logits into accessibility profiles, the profile logits were softmaxed and scaled by the exponentiated predicted log counts; and model interpretation with respect to the counts output was performed using DeepLIFT. Next, the effect allele was substituted into the sequence at the variant position, and predictions and contribution scores were obtained as for the reference sequence. For each model, we computed the variant effect as the sum of differences in per-base predicted read counts in the 100 bp window centered at variant, and computed the mean effect score across folds. We also computed the log2 fold change between predicted counts for the effect versus the non effect allele for the peak region, where a log2 fold change > 0 indicates the effect allele was predicted to increase accessibility. In figures, the mean predicted profiles and contribution scores across folds are shown.

---

**Fig. S1:**
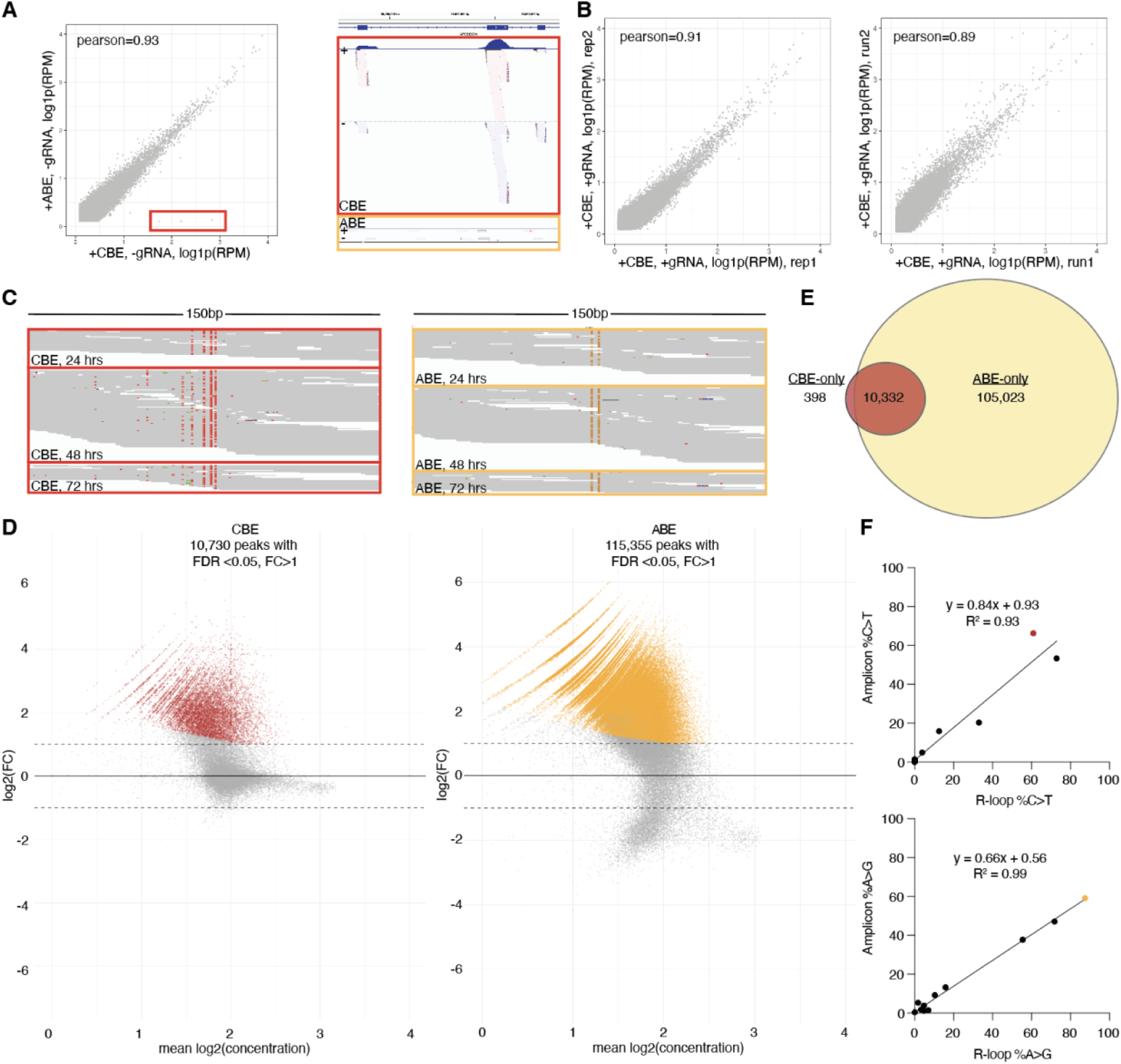
HEK293T beCasKAS. **A)** Outlier ssDNA signal highly enriched in +CBE, -gRNA conditions and not +ABE, -gRNA conditions. IGV screenshot showing the endogenous APOBEC3A gene. **B)** Correlation of +CBE, +gRNA conditions from replicate experiments performed on the same day (left) or on independent days (right). **C)** Same representative off-target peak as in Fig. 1 showing difference in beCasKAS reads on different days. **D)** DESeq2 generated MA-plots for discovering gRNA dependent R-loops for CBE +/- gRNA (left) and ABE +/- gRNA (right). **E)** Overlap of DESeq2 called peaks for CBE and ABE (FDR<0.05, FC>1). **F)** Correlation of allele fractions for amplicon sequencing for beCasKAS sequencing reads.

**Fig. S2:**
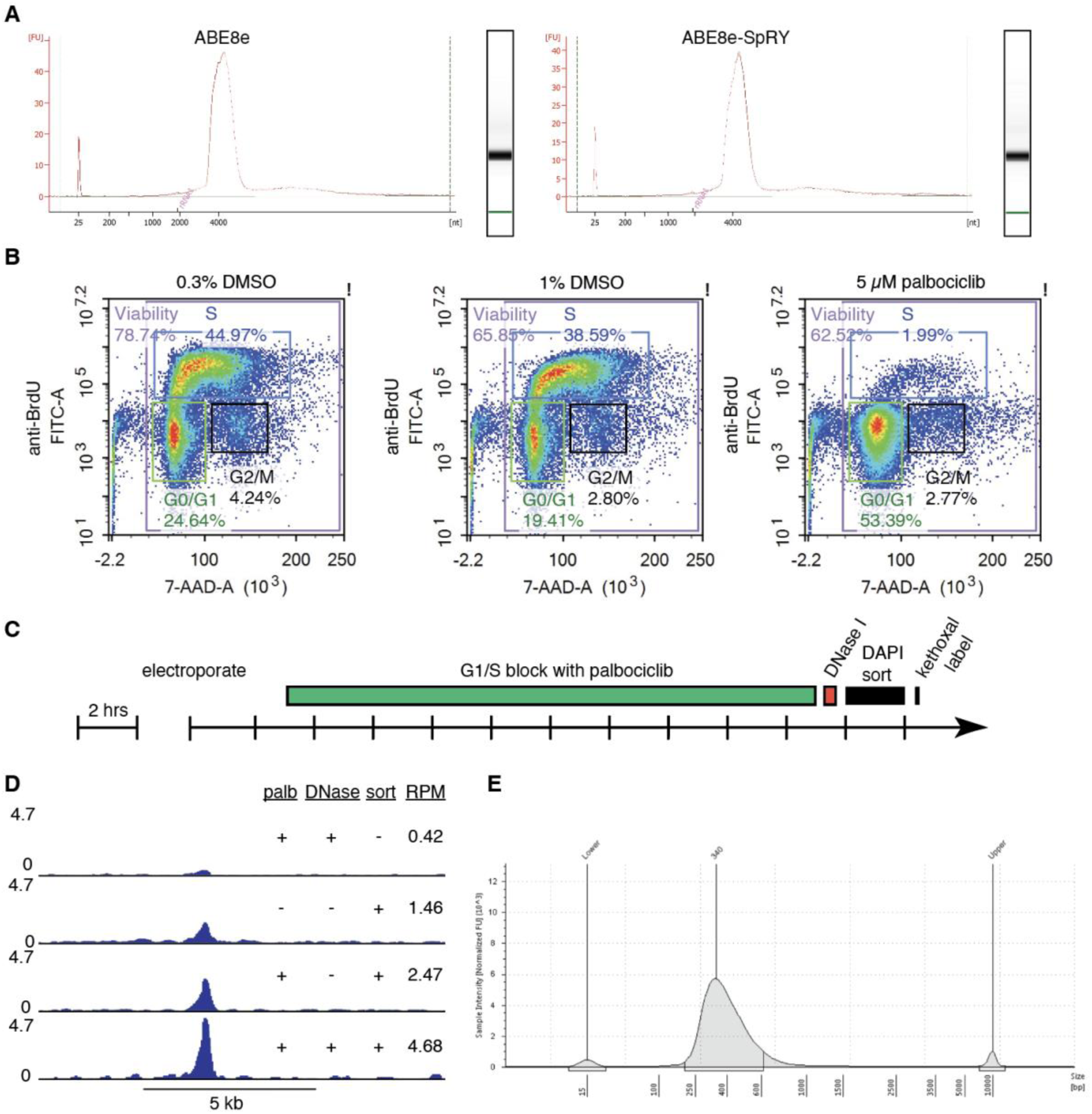
Primary human T cell beCasKAS quality control and optimization. **A)** BioAnalyzer traces verifying purity of mRNAs used in this study. **B)** Cell cycle plots of activated T-cells treated overnight with DMSO (vehicle) or Palbociclib. **C)** Primary cell beCasKAS workflow. **D)** HEK4 on-target site against three independent variables: palbociclib G1 block, DNase I treatment, and DAPI viability sorting, which all improve ssDNA signal at on-target site. The peak RPM value is provided. **E)** Example final beCasKAS library.

**Fig. S3:**
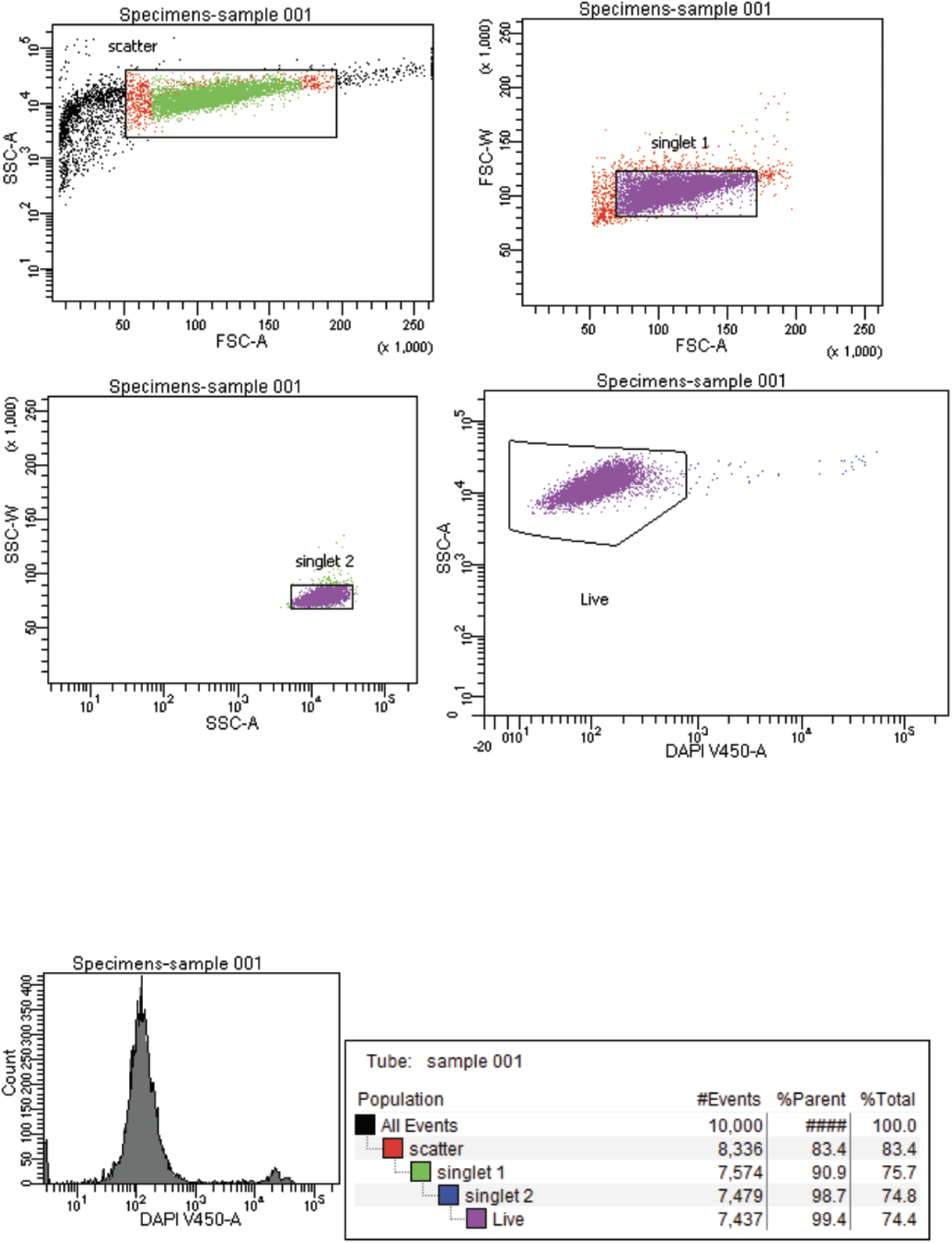
**Gating strategy for viable singlet T-cells**

**Fig. S4:**
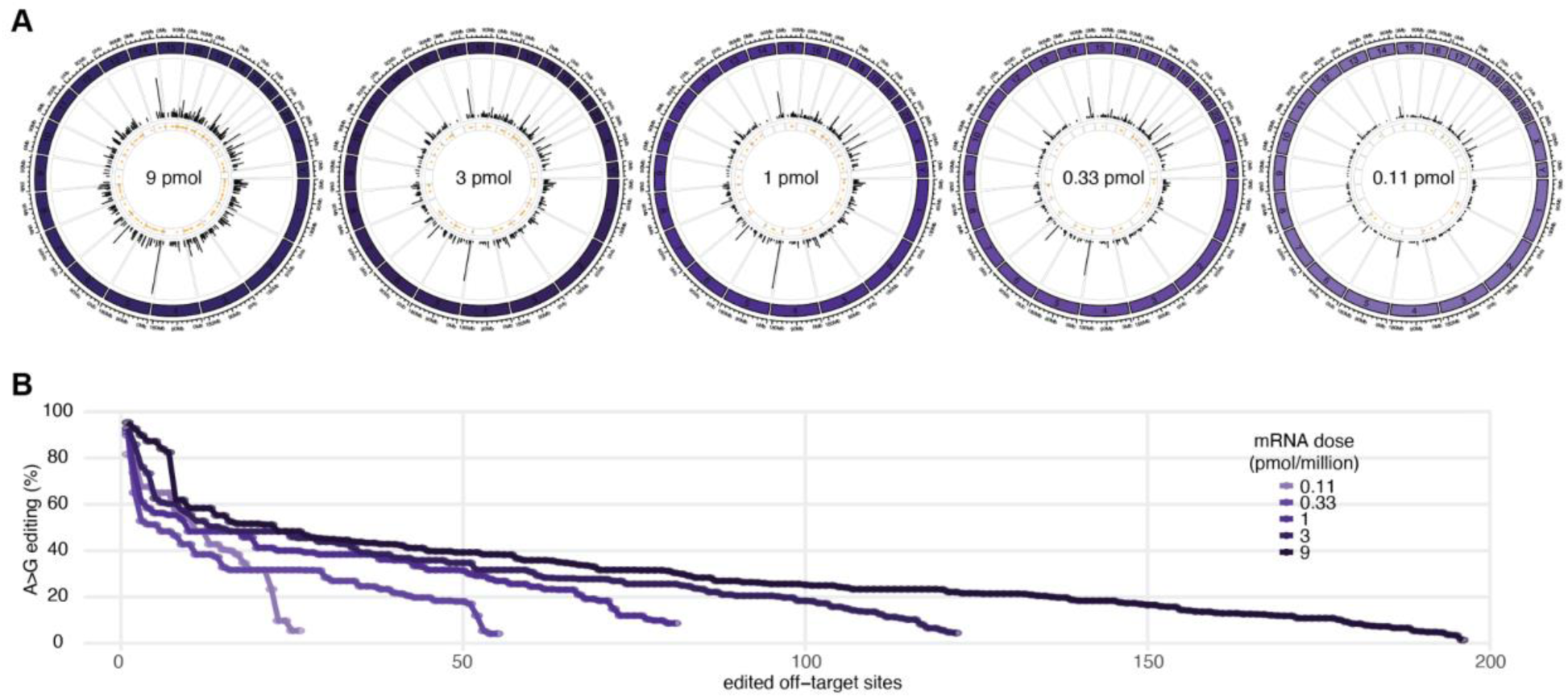
ABE8e off-target sites at different mRNA doses (pmol/million T cells). **A)** 339 R-loops (black lines) and edited sites (yellow dots) genome-wide. **B)** Editing frequency at sites within each off-target R-loops.

**Fig. S5:**
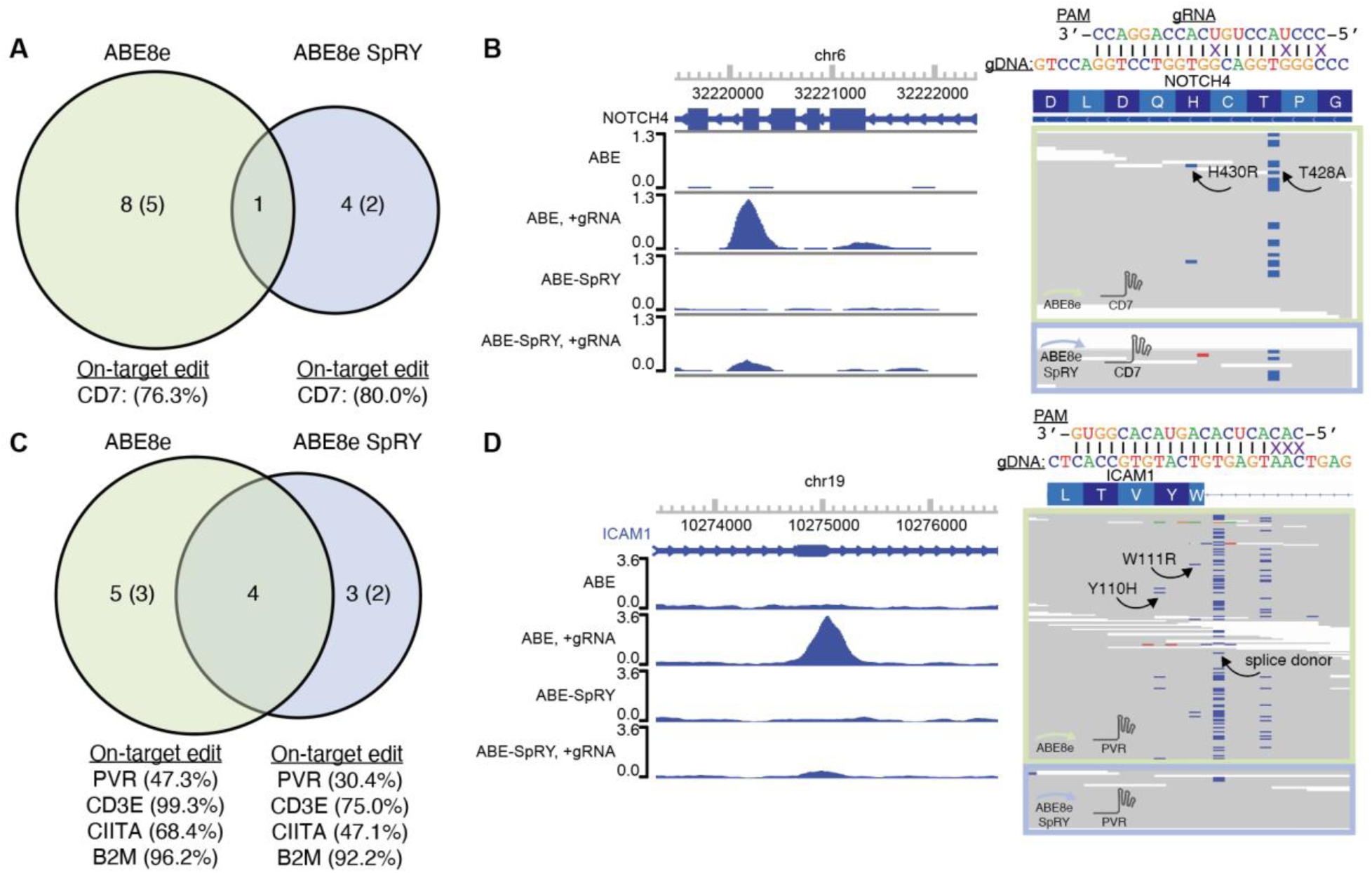
Off-target sites for therapeutic targets in primary human T-cells. **A)** Off-target R-loops for CD7 splice site targeting gRNA. The number of edited R-loops is shown in parentheses. **B)** Example off-target site in NOTCH4 tumor-suppressor/oncogene. **C)** Off-target R-loops for quadruple edited T-cells. The number of edited R-loops is shown in parentheses. **D)** Example off-target site disrupting ICAM-1 splice donor.

**Fig. S6:**
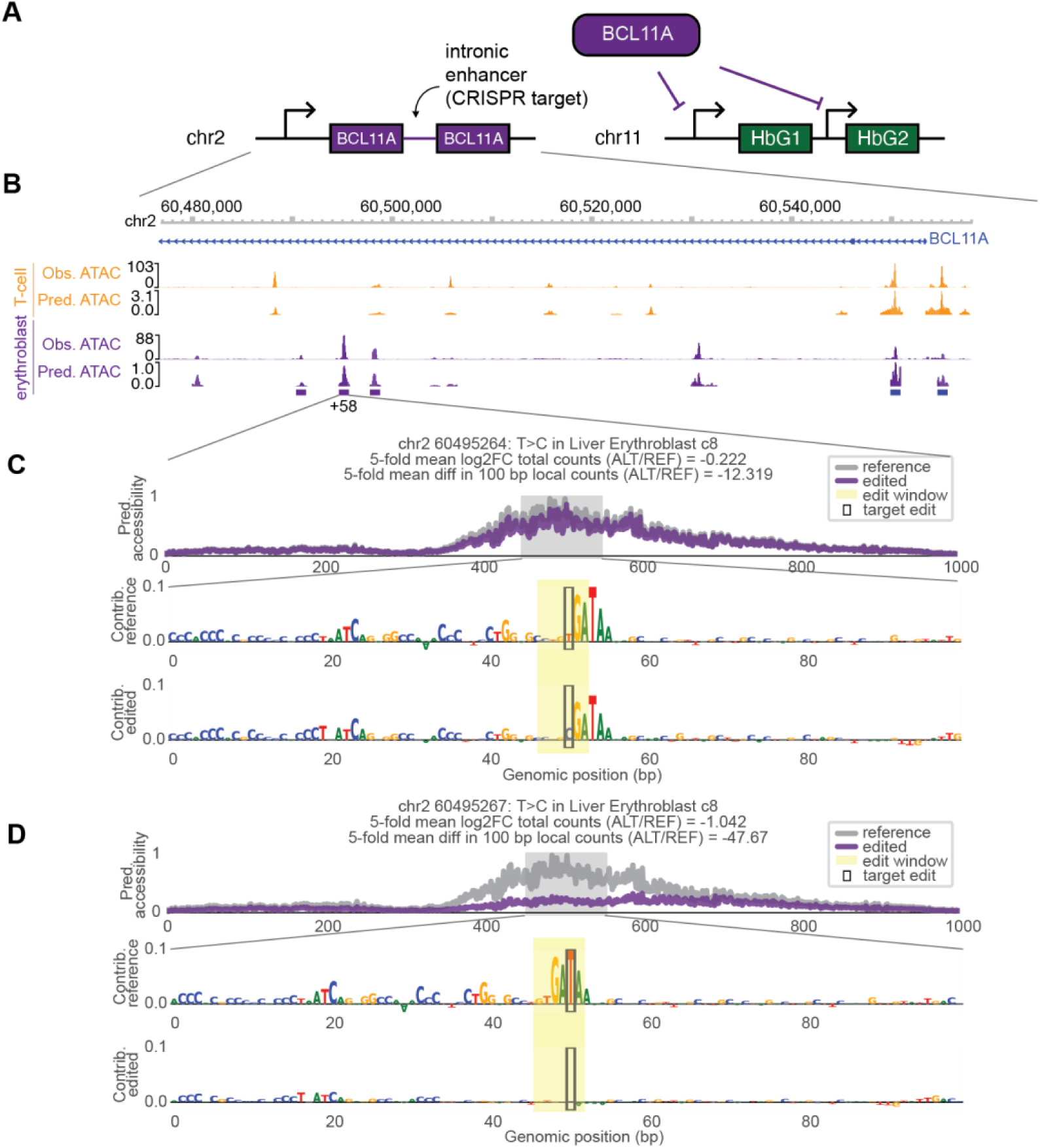
Erythroblast ChromBPNet model predicts impact of known therapeutic CRISPR target (exagamglogene autotemcel, Casgevy) on accessibility. **A)** Schematic of Casgevy mechanism. BCL11A represses fetal hemoglobin in adulthood. Disruption of a BCL11A enhancer reactivates fetal hemoglobin. **B)** Observed ATAC and predicted accessibility from ChromBPNet BCL11A intron, for T-cells (this study) and erythroblasts (*48*). **C)** Predicted accessibility for chr2 60495264: T>C edit and reference sequence in erythroblasts, along with DeepLIFT contribution scores. **D)** Predicted accessibility of chr2 60495267: T>C edit and reference sequence in erythroblasts. The ABE8e edit window is derived from the highest efficiency gRNA sg1620 (*51*).

**Fig. S7:**
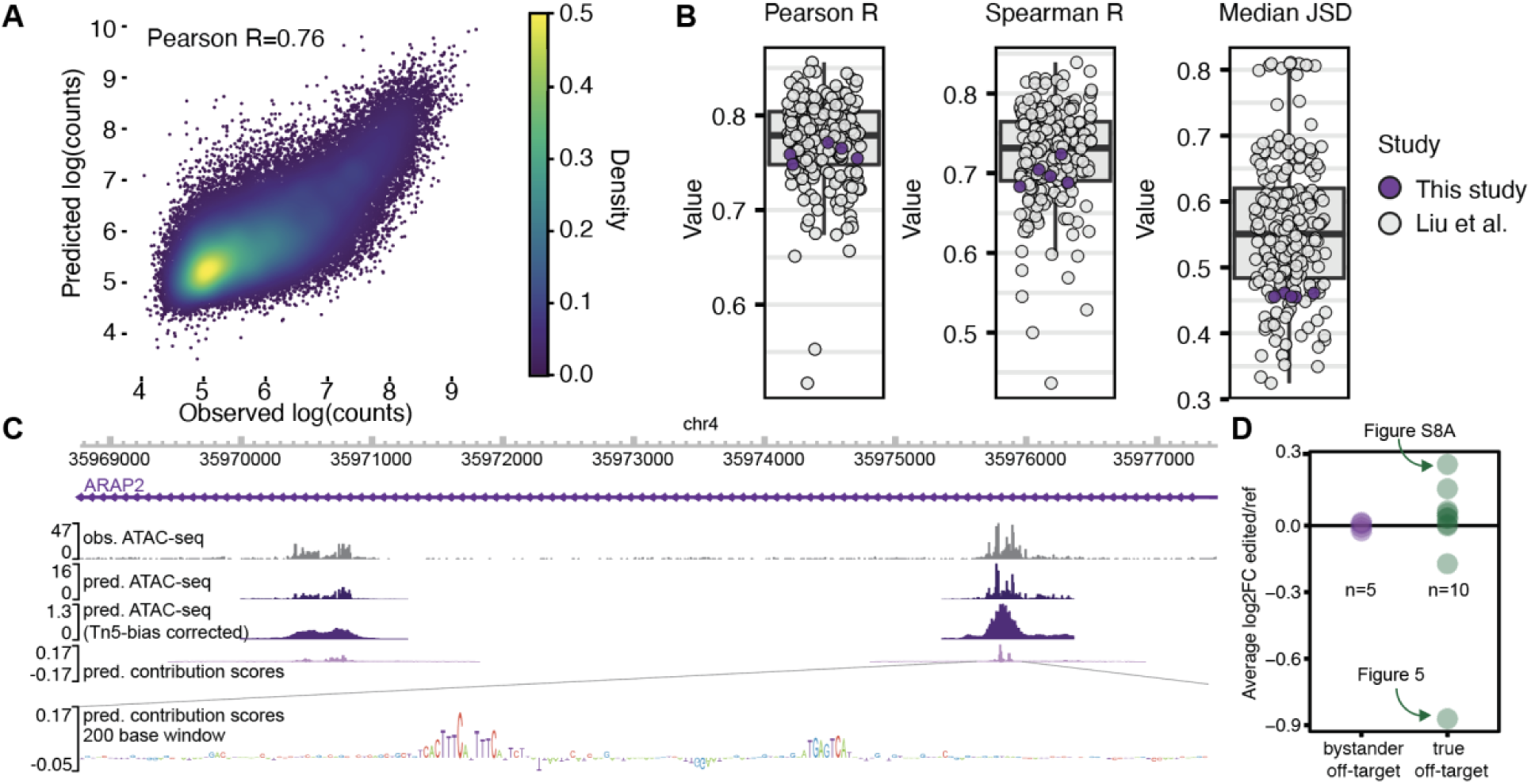
ChromBPNet model trained on primary human T-cells accurately predicts chromatin accessibility. **A)** Correlation of observed ATAC-seq log fragment counts in and predicted log counts from ChromBPNet, in peak regions, for one model fold. **B)** Performance of 5-fold models in this study compared to 189 models from prior study (*48*). Higher R and Lower Jensen-Shannon Distances (JSD) are more favorable. **C)** Example browser track showing observed accessibility (ATAC-seq Tn5 insertions), predicted accessibility from ChromBPNet, and Tn5 bias-corrected predictions from ChromBPNet. **D)** Log2 fold-changes of ChromBPNet-predicted accessibility for edited vs reference sequences by *in silico* mutagenesis, averaged over 5-fold models. Each point represents a unique off-target edit, and edits are stratified into bystander edits (occurring within the on-target R-loop) and true off-targets (occurring in an off-target R-loop).

**Fig. S8:**
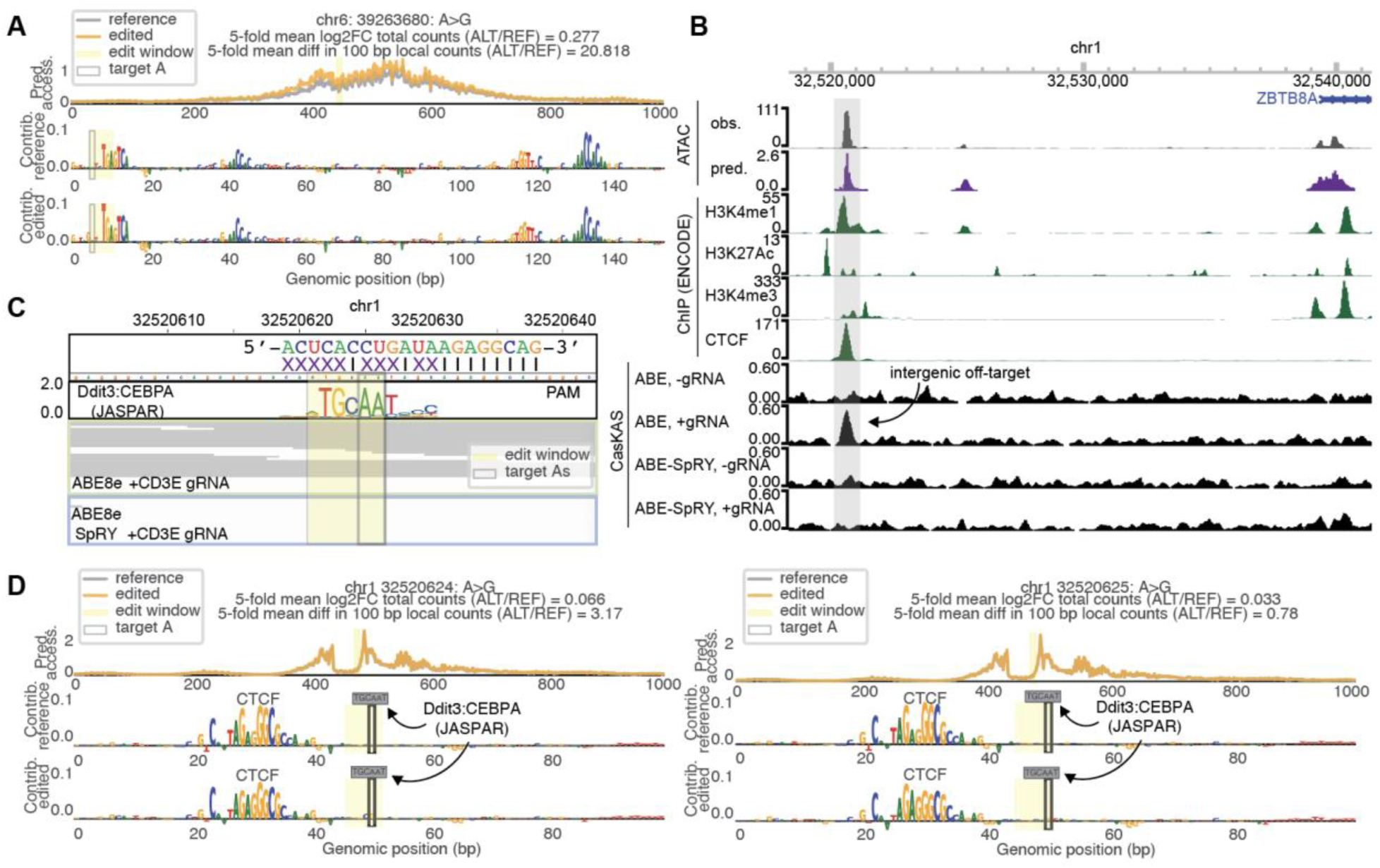
Additional ChromBPNet *in silico* mutagenesis examples. **A)** Predicted effect of chr6: 39263680: A>G mutation on accessibility. **B)** Off-target ABE8e + CD3E gRNA site overlapping with ATAC, H3K4me1 ChIP-seq, and CTCF ChIP-seq. **C)** Putative gRNA binding site with eight consecutive seed matches (black lines) and canonical GGG PAM. PWM for JASPAR-identified motif based on *in vitro* SELEX data. **D)** Predicted effect of chr1 32520624: A>G and chr1 32520625: A>G mutations on accessibility. The neighboring motif was identified as a CTCF motif by TOMTOM (p-value 2.49x10^-4^).

**Fig. S9:**
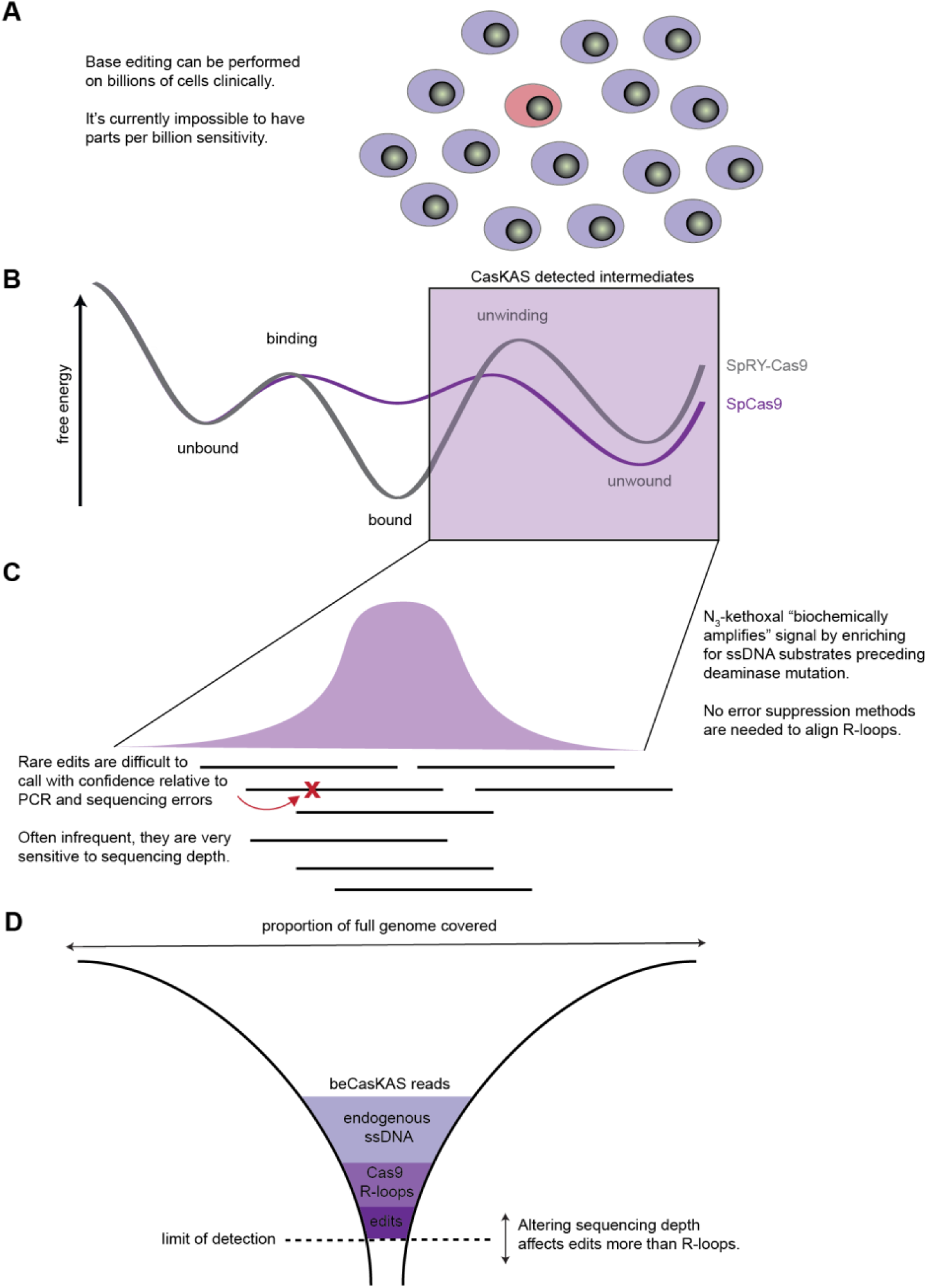
Signal amplification in beCasKAS. **A)** Scale of common clinical base editing workflows. **B)** Cartoon free energy diagram of SpCas9 and SpRY-Cas9 adapted from ref (*43*). **C)** Example gRNA-dependent beCasKAS peak. **D)** Breakdown of reads sequenced by beCasKAS.

**Table S1:**
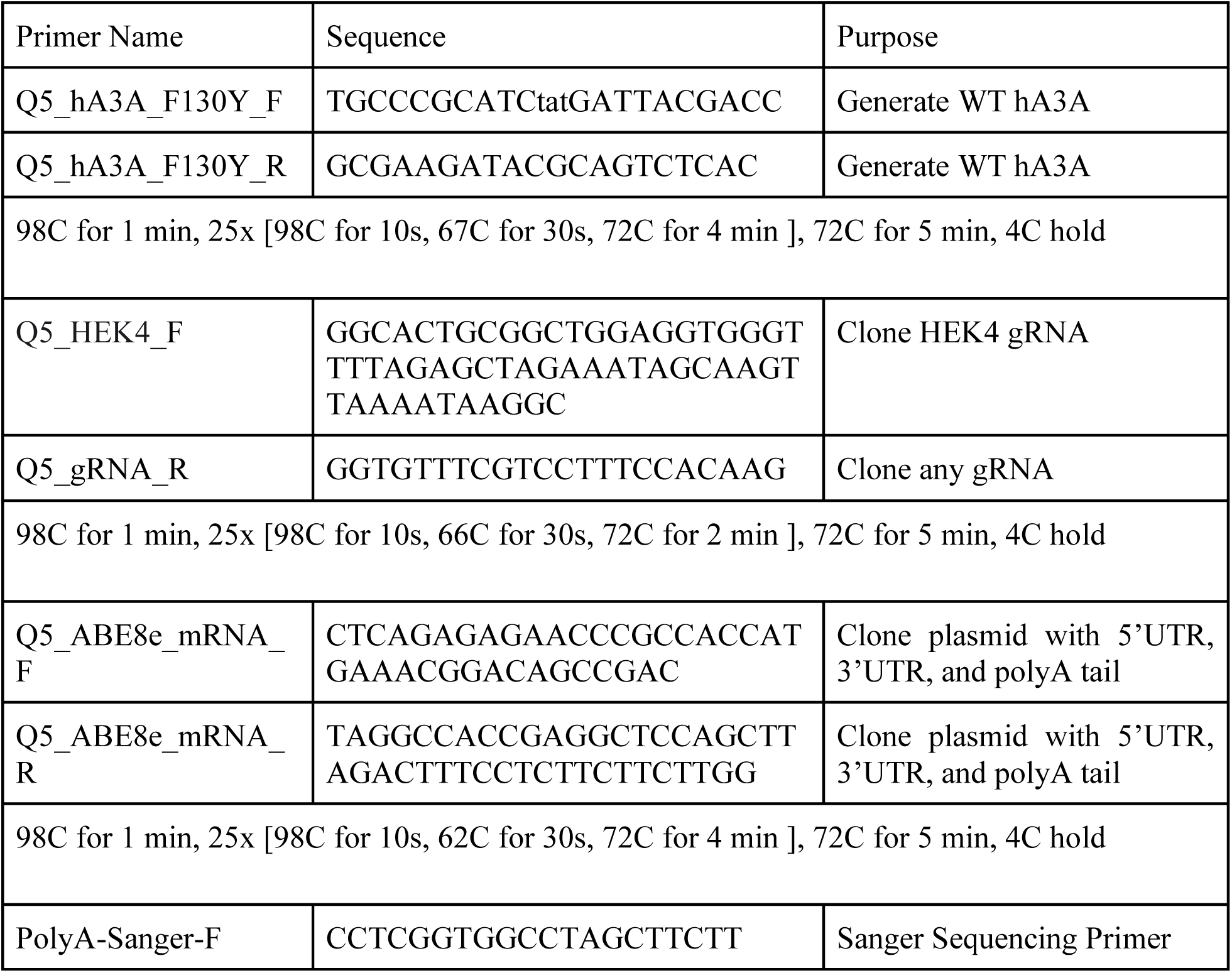
Primers and PCR conditions.

**Table S2:**
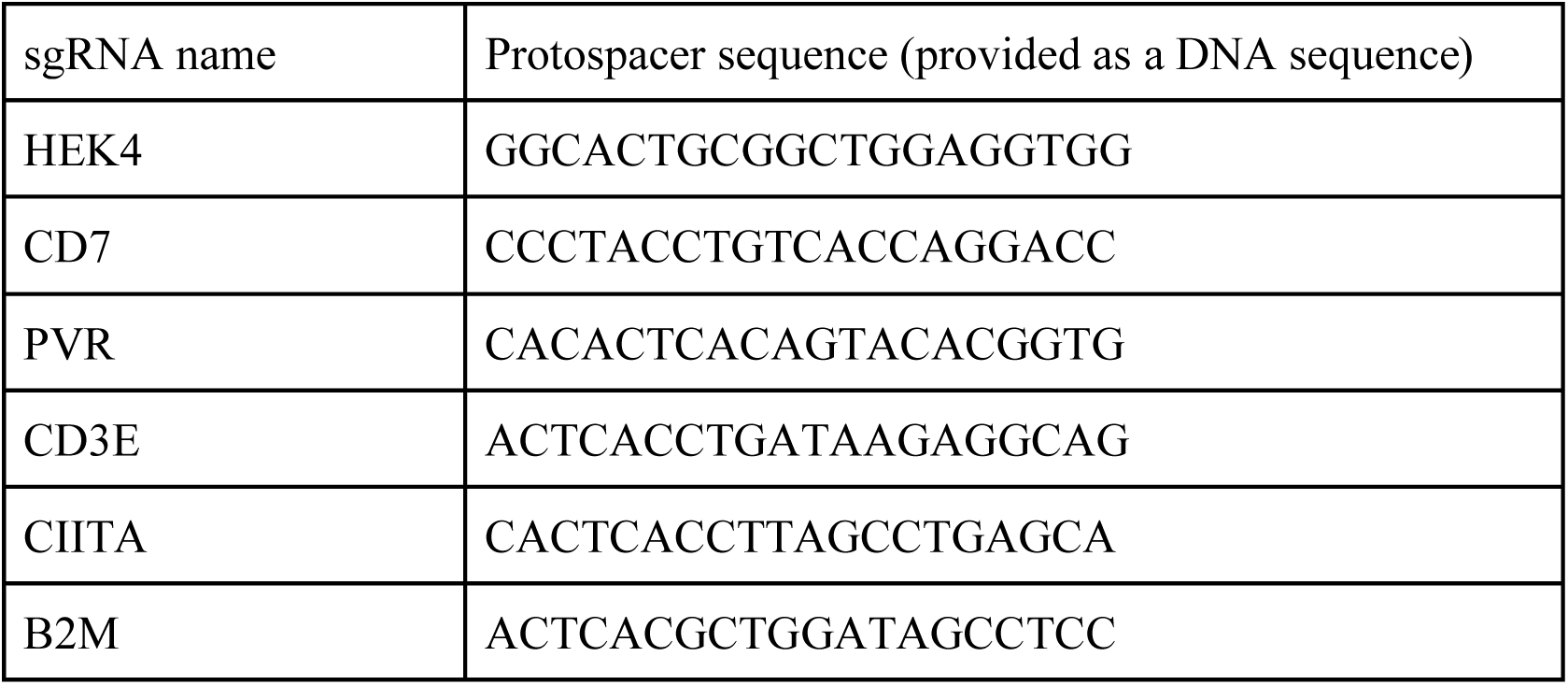
sgRNAs.

**Table S3:** ChromBPNet performance metrics.

| Fold | Spearman correlation<br>(predicted vs<br>observed log<br>counts) | Pearson correlation<br>(predicted vs<br>observed log<br>counts) | Median Jensen-Shannon distance<br>(predicted vs<br>observed profile) | Max Tn5 response |
| --- | --- | --- | --- | --- |
| 0 | 0.696 | 0.759 | 0.455 | 0.001 |
| 1 | 0.688 | 0.754 | 0.461 | 0.001 |
| 2 | 0.704 | 0.765 | 0.456 | 0.001 |
| 3 | 0.683 | 0.748 | 0.461 | 0.001 |
| 4 | 0.724 | 0.771 | 0.456 | 0.001 |
| mean | 0.699 | 0.760 | 0.458 |  |

